# The role of charge, hydrophobicity, and cooperativity in target search of SOX2 and ESRRB

**DOI:** 10.64898/2026.02.17.706345

**Authors:** L. Vanzan, C. Deluz, L. Font, D.M. Suter

**Affiliations:** Ecole Polytechnique Fédérale de Lausanne, School of Life Sciences, Institute of Bioengineering, Lausanne, Switzerland

## Abstract

Transcription factors (TFs) locate and bind specific DNA sequences within a crowded nuclear environment, yet how their biophysical properties and interactions influence their search remains unclear. Here, we modulate hydrophobicity and charge of the pluripotency TFs SOX2 and ESRRB, and combine single-molecule imaging with genomic analyses to dissect their target search dynamics. We show that increased hydrophobicity impairs specific binding while promoting confined intranuclear interactions, whereas additional negative charges decrease sampling of weak binding sites. While SOX2 retains effective target site recognition despite decreased search efficiency, impairment of ESRRB search results in a global loss of genome occupancy and redirection to SOX2-bound regions, indicating a strong reliance of ESRRB on TF-TF interactions for locating its binding sites. Single molecule imaging analysis of ESRRB search upon SOX2 depletion demonstrated that SOX2 reduces ESRRB search time by restricting its nuclear diffusion and facilitating stable DNA binding. Together, our findings highlight how TF biophysical features and cooperativity jointly shape the efficiency and specificity of target search in the nucleus.

## Introduction

Transcription factors (TFs) are sequence-specific DNA-binding proteins that regulate gene expression by locating and stably binding defined DNA motifs within gene regulatory regions ^1^. How TFs locate their target sites in the crowded and dynamic nuclear environment with sufficient speed and specificity remains a long-standing question in molecular biology. The facilitated diffusion model states that TFs alternate between three-dimensional diffusion and one-dimensional sliding or hopping along DNA to accelerate their search ^2–4^. While this model can quantitatively explain prokaryotic search ^5–8^, whether it recapitulates TF search on chromatinized eukaryotic DNA is less clear. Evidence from studies in the last few years point to a “guided exploration” mechanism, in which eukaryotic TFs switch between fast diffusion in the interchromatin compartment and slow confined diffusion within dense chromatin regions. In this model, interactions between protein and nucleosomes contribute to the confinement and to the local, repeated sampling of sites ^9–12^. Binding to functional sites is ultimately achieved through specific hydrogen bonds and van der Waals interactions formed between DNA bases and amino acids within the DNA-binding domain (DBD), which defines sequence specificity ^1^. On the other hand, the transient, non-specific and short-lived interactions with low affinity sites that accelerate target search are largely mediated by intrinsically disordered regions (IDRs) outside of the DBD ^10,12–15^. While lacking sequence conservation, IDRs are enriched in charged, polar, or low-complexity amino acids ^15,16^. Transactivation domains (TADs), which are often acidic and hydrophobic ^17,18^ are generally embedded in IDRs. Single molecule imaging and kinetic experiments showed that IDRs impact diffusion rates and residence times ^14,17^ and facilitate TF confinement into mesoscale compartments ^12,17^. However, IDR-swapping experiments ^13,19–21^, do not allow to unambiguously distinguish the contribution of IDR chemical properties such as charge and hydrophobicity from their interactions with co-activators ^18,22,23^ and nucleosomes ^24,25^ controlling target selectivity ^13,26^. In addition, post-translational modifications may further reshape TFs by altering charge, conformation, or protein-protein interactions, with potential effects on both search and binding. Phosphorylation, for example, introduces additional negative charges while slightly reducing hydrophobicity. It was shown that OCT4 phosphorylation reduces its ability to engage closed chromatin and impairs pioneer activity (Shin et al., 2016), while phosphorylation of FOXA1 increase its interphase mobility, consistent with a looser chromatin engagement during search (Zhang et al., 2024).

In eukaryotes, TF binding is also modulated by cooperativity with other TFs, which enables combinatorial logic in enhancer function and expands the regulatory potential of TF networks ^27^. In pluripotent stem cells, cooperative interactions within the core network of SOX2, OCT4, NANOG, ESRRB, are critical for maintaining regulatory element activity ^28^. SOX2 can redirect OCT4 and NANOG to alternative loci ^29^ and depletion of ESRRB or NR5A2 diminishes the occupancy of SOX2, OCT4 and NANOG at key regulatory elements ^30^. Recent studies have shown that the formation of TF homodimers or heterodimers extends residence times ^31–35^, slows down nuclear diffusion ^14^ and allows occupancy at low affinity sites ^36^ or at composite motifs ^27,37^. Mechanistic and modeling works showed that ^38^ cooperative TF-TF interactions may facilitate target search to colocalized motifs ^39,40^. Studies in live cells highlighted how clustering of TF molecules at target loci accelerate search ^9,41^. However, to which extent TFs modify their target search strategies in presence or absence of other TFs is unclear.

Here, we study how electrostatic interactions, hydrophobicity and cooperativity shape the target search of two different TFs. We focus on SOX2 and ESRRB, two pluripotency factors that share overlapping target regions but differ in DBD class and in their biophysical properties. Our findings highlight how TF-specific biophysical properties and cooperative interactions converge to orchestrate efficient genome exploration and target binding.

## Results

### Characterization of SOX2 and ESRRB charge and hydrophobicity variants

We designed constructs in which Esrrb and Sox2 are fused to both a Halo-Tag and a HA-Tag (Figure 1a). We then fused a sequence encoding 18 hydrophobic amino acids to the C-terminus of SOX2 and to the N-terminus of ESRRB to generate the SOX2 and ESRRB hydrophobic variants (SOX2hydro and ESRRBhydro), and a sequence encoding 18 negatively charged amino acids to SOX2 to create the SOX2 negative variant (SOX2neg) (Figure 1A - C). We also generated an ESRRBneg variant, but unfortunately we were unable to express it at detectable levels. Adding the Halo-Tag and charged/hydrophobic residues at opposite ends of the coding sequence allows minimizing the risk of altering Halo dye photophysics by the added residues. We measured the average hydrophobicity and net charge of these TF variants (vTFs) (Figure 1B-C) (see Methods) and verified that they were comparable to those of the other members of their respective TF families (Figure S1A-B). We then modelled the 3D structure of vTFs and predicted their DNA binding specificity using Alphafold 3 ^42^. We found that the addition of hydrophobic or negatively charged amino acids did not visibly alter the folding of either SOX2 or ESRRB or their predicted ability to bind DNA (Figure S1C-G). We then integrated the wt and vTFs fusion constructs by lentiviral transduction into the genome of mouse ESCs expressing the rtTA3G transactivator, allowing for controlled vTF expression by adjusting doxycycline (dox) concentrations (Figure 1D-E). We generated monoclonal cell lines with comparable vTF expression levels upon dox induction as measured by Halo imaging (Figure S2A). We then checked the binding profiles of Halo-HA–tagged and endogenously-expressed, wt forms of ESRRB and SOX2 by ChIP-seq, and found these to be comparable (Figure S2B-E). This indicates that the Halo and HA tags did not interfere with the ability of SOX2 and ESRRB to engage their genomic targets.

**Figure 1.**
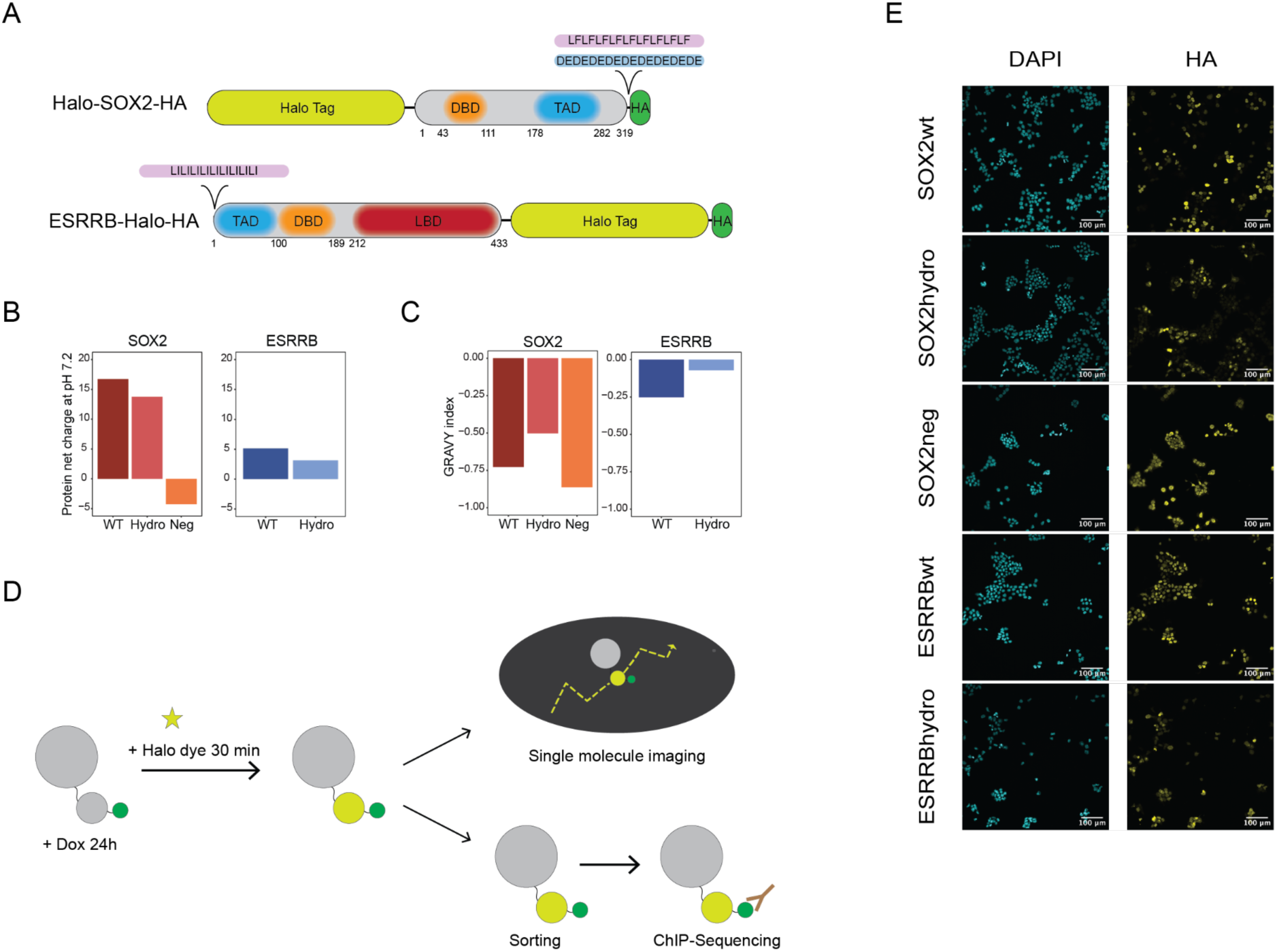
Generation of TF variants fusion constructs. **A**) Mouse SOX2 (top) and ESRRB (bottom) cDNAs were fused to a Halo-tag for fluorescence labeling, and with a HA-tag for chromatin immunoprecipitation (ChIP). 18 hydrophobic (purple) amino acids were added to SOX2 and ESRRB to generate SOX2hydro and ESRRBhydro, and 18 negatively charged (blue) amino acids were added to SOX2 to generate SOX2neg. **B**) Protein net-charges for SOX2 (left) and ESRRB (right) WT and variants at pH 7.2. **C**) GRAVY (Grand Average of Hydropathy) index of hydrophobicity calculated for SOX2 (left) and ESRRB (right) WT and variants using the Kyte & Doolittle scale ^43^, where positive values indicate higher hydrophobicity (see Methods). **D**) vTFs fusion constructs were cloned downstream of a TRE3G promoter and transduced into CGR8 mouse ESCs expressing the rtTA3G transactivator. Upon dox addition, rtTA3G binds to the TRE3G promoter, inducing expression of the vTF. Fusion proteins were visualized upon Halo staining. vTFs were expressed at low concentrations and sparsely labelled with Halo-dyes for single molecule imaging experiments. To study genomic binding, cells were sorted based on Halo fluorescence and ChIP-Seq was performed using an anti-HA antibody. **E**) DAPI and HA-Tag immunostaining of vTF cell lines after 24h of dox induction.

### TF hydrophobicity promotes slow and constrained diffusion

We measured the diffusive behaviours of our vTFs using fast single molecule tracking (SMT). To reach single molecule resolution, we expressed vTFs using low dox concentrations (10-25 ng/µL depending on the variant), and labelled them using picomolar Halo dye concentrations. We recorded 100Hz movies at high laser power for 1 min using Highly inclined thin illumination (HiLo) microscopy (Figure 2A-B and Supplementary Table 2). We extracted single molecule trajectories using TrackIT ^44^ and generated cumulative jump distance distributions that we fitted with a three-component Brownian diffusion model, as previously reported to be the best fit for SOX2 ^19,45^ and other TFs ^11,46–49^ (Figures 2B, S3A-B and Supplementary Tables 3). This model describes quasi-immobile, slow and fast diffusing populations corresponding to bound molecules and molecules diffusing in regions of high and low DNA density, respectively ^11^. The bound population includes both specific and transient binding events. Binding was not saturated in our overexpression conditions, as evidenced by the linear relationship between bound and total detected molecules ^49^ (Figure S4A). Analysis of diffusion fractions (Figure 2C) and diffusion coefficients (Figure S4B) revealed partially different behaviours for SOX2wt and ESRRBwt: while bound fractions were similar (∼ 28%), SOX2wt showed a preference for fast diffusion, as previously reported ^45^, while ESRRBwt favoured slow diffusion. This suggests that these two TFs might employ different search strategies. Hydrophobic variants of SOX2 and ESRRB displayed increased fractions of bound and slow-diffusing molecules, and a marked decrease in the fraction of fast-diffusing ones. In addition, the hydrophobic variants were markedly slower than their respective WT counterparts in the bound and slow diffusing fractions, while showing accelerated diffusion speed in the fast component (Figure S4B). SOX2neg displayed a reduced bound fraction, and an increase in both slow and fast diffusing fractions, with diffusion speeds unchanged (Figure S4B). Overall, these results suggest that hydrophobicity slows down TF mobility. We confirmed our finding by calculating the average diffusion coefficient D_eff_ as a measure of the overall mobility ^44^, which was indeed decreased for the hydrophobic variants but not for SOX2neg (Figure 2D). We observed a significant linear negative correlation between D_eff_ and hydrophobicity, while more charged proteins (both positively and negatively) tended to increase D_eff_ (Figure S4C and methods). Next, we asked if hydrophobicity may impact confinement of diffusing molecules. We used a Hidden Markov Model approach ^50^ to classify trajectory segments as bound or free (Figure 2B). We filtered out bound segments to avoid measurement bias, and verified that the residual bound fraction was below 1% ^11^ (Figure S4D and Methods). To assess diffusion anisotropy, we measured the angles formed between consecutive displacement vectors along single-molecule trajectories, and plotted the angle distribution probability. A peak at 180° in the angle distribution reflects a bias toward backward motion. Both SOX2hydro and ESRRBhydro exhibited a markedly sharper peak at 180° (Figure 2E), as also quantified by the “fold anisotropy” ratio (*f_180/0_*) ^10^, which captures the relative probability of backward versus forward displacements. (Figure 2F). When *f_180/0_* was analyzed as a function of mean displacement length, SOX2wt and SOX2neg both peaked at 100–300 nm before transitioning to more isotropic diffusion over larger distances, in line with previously reported compact nuclear environment exploration before shifting to more isotropic diffusion^10–12^. SOX2hydro followed a similar pattern, but with much stronger anisotropy, indicating enhanced confinement (Figure 2G). ESRRBwt showed a relatively constant *f_180/0_* across all displacement lengths, indicating that ESRRBwt diffusion behaviour is uniform at all distances. ESRRBhydro showed very high anisotropy at shorter displacements (up to ∼250 nm) before decreasing at larger distances (Figure 2G). Together, these results suggest that hydrophobicity constrains diffusion but its effect becomes less pronounced over longer distances, possibly reflecting transitions between local and more global modes of nuclear exploration. Note that while increased confinement could be mediated through interactions with chromatin, we cannot exclude that interactions with other nuclear components may also contribute. The high anisotropy at short distances might also be due to persistent trapping interactions making hydrophobic variant TFs unable to escape their local environment.

**Figure 2.**
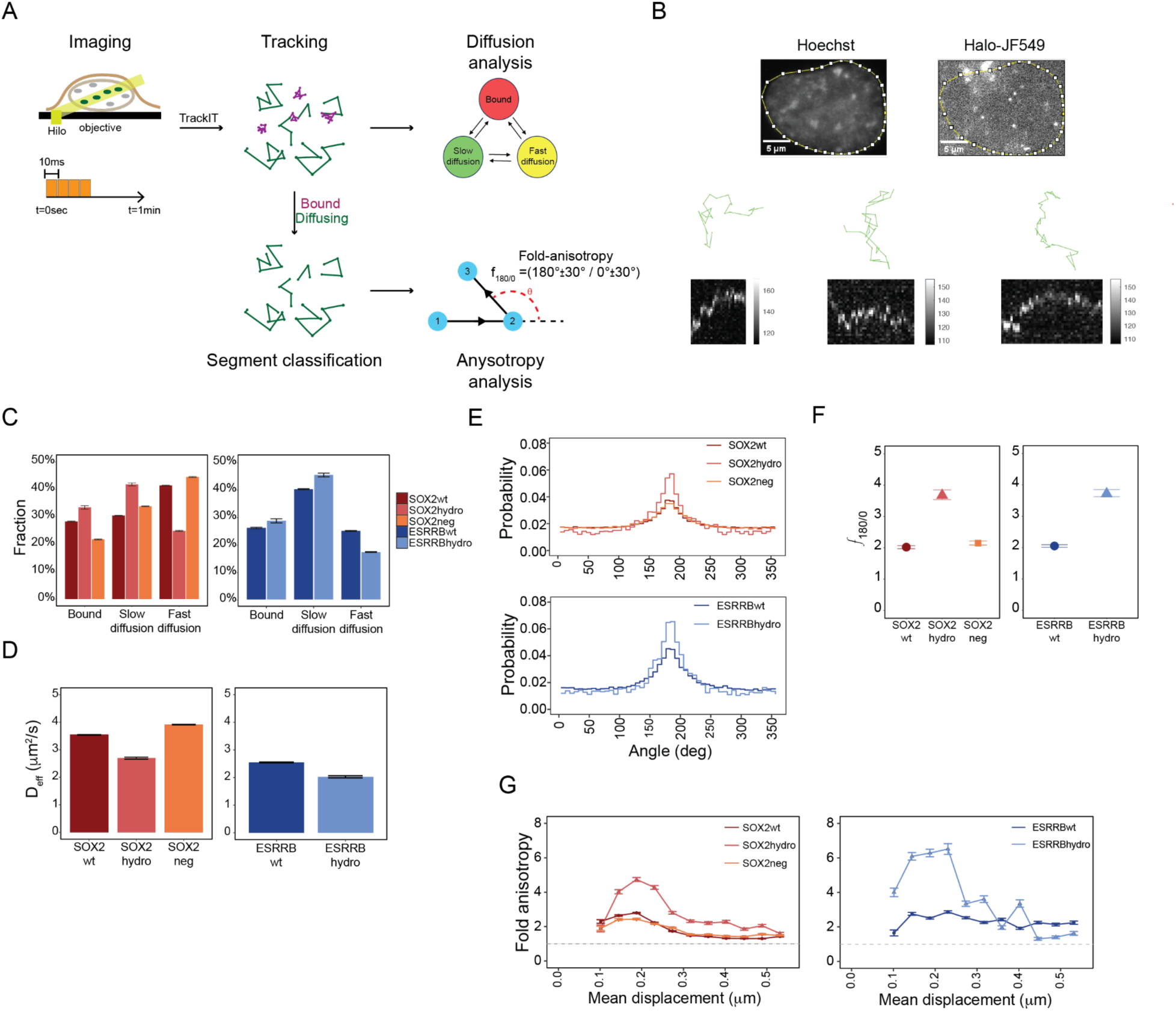
Addition of hydrophobic or negatively-charged residues alter TF diffusion and anisotropy. **A**) Schematic diagram of illumination pattern for continuous diffusion movies (with indicated camera integration time, frame cycle time and total imaging time) and of the analysis pipeline. **B**) (top) Representative images of Hoechst-labelled nuclei and sparsely labelled JF549-labelled Halo-TF molecules; (bottom) trajectories and kymographs of representative diffusing molecules. **C**) Fractions of bound, slow and fast diffusing molecules, derived from fitting continuous movies with a three-states diffusion model (Methods and Supplementary Table 3). Data are provided as mean values ± standard deviation of 500 resamplings with 80% of the data. **D**) Average diffusion coefficient D_eff_ as measure of overall TF mobility, derived from bound fractions in Figure 2c and diffusion coefficients in Figure S4b. Data are provided as mean values ± standard deviation of 500 resamplings with 80% of data. **E**) Histograms of angle distribution probability for wt and variant TFs. **F**) Fold anisotropy ratio *f_180/0_*. Data are presented as mean values ± standard deviation. **G**) Fold anisotropy ratio *f_180/0_* as function of mean displacement length over all lag times. Data are provided as mean values ± standard deviation of 500 resamplings with 80% of data.

### Negative charges decrease the fraction and residence time of long binding events

We next aimed at quantifying the impact of hydrophobicity and negative charges on TF dissociation rates. We employed several time-lapse HILO conditions, keeping a fixed camera integration time of 60ms and variable dark times, to obtain frame cycle times ranging between 150 ms and 5 s (Figure 3A and Supplementary Table 4). This approach ensures robust tracking across an extended temporal range, while correcting for fluorophore photobleaching ^44^. We generated survival time distributions from tracked data (Figure S5A) and from them inferred dissociation rates using GRID ^51^, which applies an inverse Laplace transformation to survival time distributions to resolve populations of bound molecules. The resulting dissociation rates spectra (Figure S5B-C) capture the relative fraction of binding events and the associated residence times reflecting multiple binding states observed in the measurement window.

**Figure 3.**
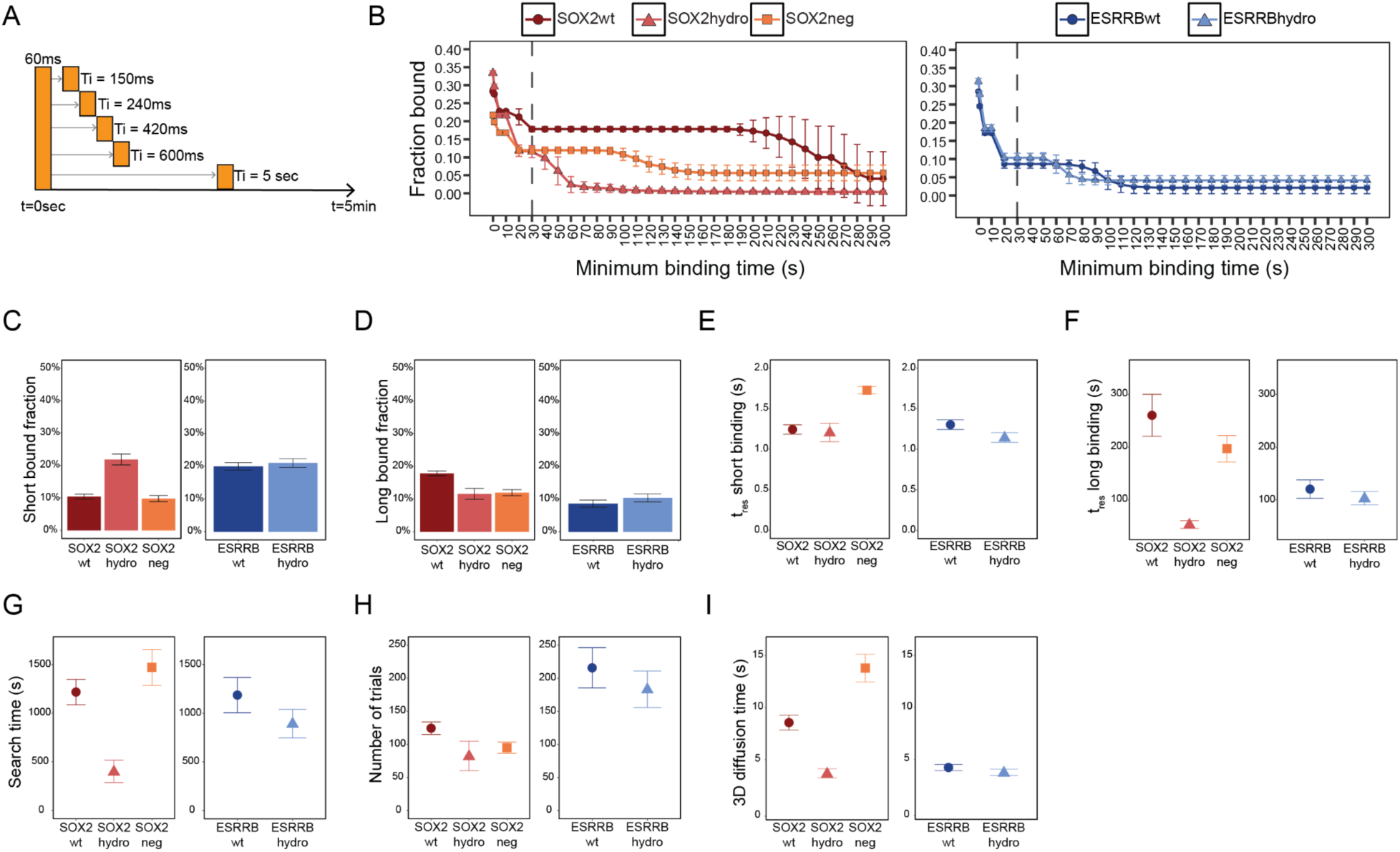
Binding and search kinetics are strongly impacted by negative/hydrophobic residues. **A**) Illumination patterns of time-lapse movies with indicated camera integration time, frame cycle times and total imaging time. **B**) Cumulative binding curves showing the proportion of molecules that remain bound for at least the indicated residence time. Curves were obtained by weighing the total bound fraction by GRID dissociation rate amplitudes and plotting it as a function of residence time (see Methods). At binding time 0 s, the corresponding value corresponds to the total bound fraction determined by continuous diffusion movies (Figure 2c and Supplementary Table 4). Data are provided as mean values ± SD of 500 resamplings with 80% of data. **C-F**) Average fraction (**C,D**) and corresponding average residence times (**E,F**) of all binding events lasting less than 30 s (**C,E**) and more than 30 s (**D,F**). Data are provided as mean values ± SD. **G**) Search time needed for a single TF molecule to find a specific site (Methods and Supplementary Table 5). **H**) Number of unspecific encounters determined for each TFs from the amplitude of the event spectrum above 30s (Methods and Supplementary Table 5). **I**) Time spent diffusing between binding events (Methods and Supplementary Table 5). For panels G, H, I data are provided as mean values ± error from error propagation.

We first wondered whether SOX2 and ESRRB molecules predominantly engage in short, transient binding or stable, potentially functional interactions, and whether the relative proportions of these interaction modes change upon addition of hydrophobic or negatively-charged residues. Frequency and stability of individual binding events can be integrated by weighting the bound fraction obtained by diffusion analysis (Figure 2C) with the amplitude of the dissociation rates spectrum (Figure S5C and Methods). This provides a quantitative measure of binding persistence over time. By plotting this cumulative bound fraction as a function of residence times we obtained a cumulative binding curve that represents the proportion of molecules that remain bound for *at least* a given duration (Figure 3B and Methods). For all TFs, we found a sharp reduction in the fraction of bound molecules between 0 and 30 s, reflecting a large population of short-lived binding events. SOX2wt exhibited the highest bound fraction and most stable binding, with more than half of its bound molecules engaging in minutes-long binding, and only a small fraction sampling sites. In contrast, only about 2% of ESRRBwt molecules were bound longer than 90 s. ESRRBhydro and SOX2hydro maintained fractions of short-binding molecules comparable to their wt counterparts. However, their curves exhibit faster declines, with SOX2hydro being virtually unbound after 110 s, while ESRRBhydro retains only a small fraction (about 4%) of bound molecules above 80 s, which persist for over 300 s. This suggests that hydrophobicity might reduce binding stability for long-lived events. While in general a smaller fraction of SOX2neg molecules were bound in comparison to SOX2wt, the decay of bound fraction as a function of binding time was slower than for SOX2hydro, with a higher fraction bound above 90 s.

To separate transient from stable binding, we fixed a time threshold of 30 s, reflecting the time point at which most curves flatten, and determined long and short bound fractions (Figure 3C-D) and their average residence times *t_res_* (Figure 3E-F). For more hydrophobic TFs (SOX2hydro, ESRRBwt and ESRRBhydro) short binding events were more frequent (about 20%) than long ones (10%), while this was the opposite for SOX2wt. SOX2neg displayed comparable small fractions (∼10%) of both short- and long-lived binding events, consistent with the higher frequencies of slow and fast diffusing molecules (Figure 2C) and overall higher mobility (Figure 2D) than SOX2wt. Interestingly, short binding events showed comparable average residence times across SOX2wt, SOX2hydro, ESRRBwt, and ESRRBhydro, with only SOX2neg exhibiting a detectable increase (Figure 3E). In contrast, long binding events (Figure 3F) differed more substantially: SOX2wt displayed the average longest residence time (260 s), exceeding that of ESRRBwt (120 s). This stability was strongly reduced in SOX2hydro (53 s) but not in SOX2neg (197 s), whereas ESRRBwt and ESRRBhydro showed little difference, indicating that the impact of hydrophobicity on residence times is more pronounced for SOX2 than ESRRB. In line with previous observations on Figure 3B, the most hydrophobic TF variants (SOX2hydro, ESRRBwt and ESRRBhydro) have shorter long residence times than SOX2wt or SOX2neg (Figure 3F). Interestingly, SOX2neg did not display an altered fraction of short binding events compared to SOX2wt, but displayed a longer residence time for short interactions and a modestly reduced residence time for long events, a pattern that is consistent with the idea that negative charges promote more persistent, nonproductive contacts while destabilizing productive, stable binding. To estimate the target search time, defined as the average time a TF requires to locate one of its specific target sites, we applied an equilibrium binding model incorporating free diffusion and specific and nonspecific binding ^52,53^. In this framework, the search time is determined by the duration and number of unspecific “trial” binding events before a TF enters into a long-bound state, which we assume to reflect binding to a specific target sequence. We additionally determined the amount of time a molecule spends diffusing between specific binding events (see Methods). Overall, SOX2wt and ESRRBwt needed the same time to find a specific target (Figure 3G) even though SOX2wt sampled less sites than ESRRBwt and spent more time diffusing between trials (Figure 3H-I). On the other hand, ESRRBwt high number of trials is consistent with a preference for slow diffusion in a crowded nuclear environment (Figure 2C) where contacts are more probable ^11,54^. For both SOX2hydro and ESRRBhydro, total search time, diffusion time and number of trials are reduced, suggesting that high hydrophobicity might facilitate more efficient scanning by weakening long-lived interactions and thus promoting faster TF release for a new round of search. The impact of adding hydrophobic residues appears more pronounced for SOX2 than for ESRRB, which is consistent with the fact that ESRRB wt is already intrinsically more hydrophobic (Figure 1B). By contrast, SOX2neg exhibited slower search dynamics (Figure 3G). While the number of trials was slightly reduced, diffusion time increased, in agreement with the reduced total bound fraction (Figure 2C) and lower long-bound fraction (Figure 3D). Overall, these data suggest that hydrophobicity primarily destabilizes long TF binding events, thereby accelerating target search, while the introduction of negative charges increases the residence time of transient binding, thus slowing down the overall target search process.

### SOX2 but not ESRRB retains specificity when search is impaired

We next asked how differences in vTF search efficiency translate in terms of genome-wide occupancy of specific binding sites. To eliminate the impact of TF concentration on TF occupancy ^55–57^, we stained and sorted Halo-positive fluorescent cells in the G1 phase of the cell cycle, maintaining comparable mean fluorescence intensities between wt and variant TFs (Figure 4A and 4C). We then performed chromatin immunoprecipitation followed by sequencing (ChIP–seq) using an anti-HA antibody to specifically capture HA-tagged TFs. We spiked in Drosophila chromatin as a normalization method to ensure that binding enrichment could be robustly compared across TF variants. We found that binding occupancy and number of bound regions was lower for SOX2hydro (6713 peaks) and SOX2neg (3288 peaks) as compared to SOX2wt (12815 peaks) (Figure 4B and Figure S6A). A similar trend was observed for ESRRBwt and ESRRBhydro (10036 peaks for ESRRBwt against 786 for ESRRBhydro) (Figure 4D and Figure S6B). The number of ChIP-Seq peaks only mildly correlated with their occupancy, calculated by weighing the long bound fraction derived from SMT studies with the respective residence time (Figure S6C), suggesting that a fraction of the molecules immobile for 30 s in SMT might i) be non-specifically bound to DNA, ii) reflect indirect binding not efficiently captured by ChIP-seq iii) are caused by immobilization that is not mediated by DNA, or iv) that ChIP-seq additionally captures events that are not identified as long binding events. We then investigated if the nature of genomic targets was conserved across TF variants. SOX2 variants retained a substantial fraction of common peaks, while each variant also engaged with a distinct subset of regions. In contrast, ESRRBwt and ESRRBhydro exhibited markedly different binding profiles, sharing only 62 genomic loci (Figure S6D). Interestingly, both hydrophobic variants were mostly excluded from promoter regions (Figure S6E), which tend to be more accessible than enhancers on average.

**Figure 4.**
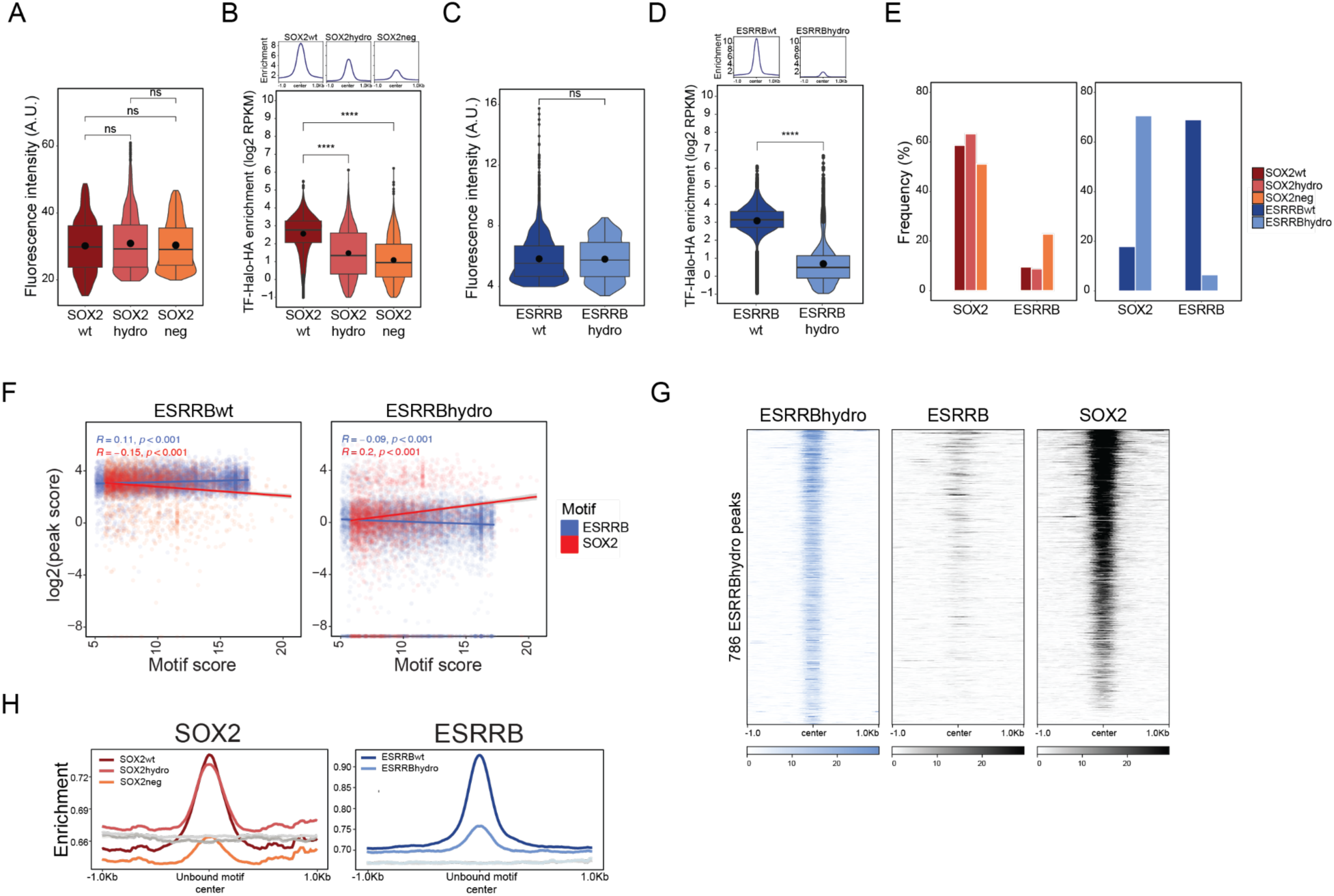
ESSRBhydro is redirected to SOX2-bound regions. **A**) Fluorescence intensity of G1-sorted SOX2wt, SOX2hydro and SOX2neg Halo-tag labelled cells (n=2 replicates). **B**) Average enrichment score (log2 RPKM) of HA ChIP-Seq in SOX2wt-, SOX2hydro- and SOX2neg-expressing sorted cells. **C**) Fluorescence intensity of G1-sorted ESRRBwt and ESRRBhydro Halo-tag labelled cells (n=2 replicates). **D**) Average enrichment score (log2 RPKM) of HA ChIP-Seq in ESRRBwt- and ESRRBhydro-expressing sorted cells. lack dots: mean. **E**) Frequency of SOX2 and ESRRB motifs at vTFs called peaks (see Methods). **F**) ChIP-Seq enrichment peak score (log2 RPKM) for ESRRBwt (left) and ESRRBhydro (right) plotted against the ESRRB or SOX2 motif quality score for the same peak. **G**) Heatmap of ChIP-Seq enrichment of ESRRBhydro, endogenous ESRRB and endogenous SOX2 at ESRRBhydro peaks. **H**) vTFs ChIP-Seq signal over all unbound SOX2 and ESRRB motifs (coloured lines) compared with background sequences without the corresponding motif (see Methods and ^60^).

We then performed motif enrichment analysis using Homer ^58^. Peaks called by variant SOX2 TFs were highly enriched in SOX2 motifs indicating that, despite the progressive reduction in overall binding occupancy, both SOX2hydro and SOX2neg retained the ability to recognize their specific target sequences. Surprisingly, while as expected the ESRRB motif was prevalent at ESRRBwt peaks, we found the SOX2 motif to be predominant at ESRRBhydro peaks (Figure 4E). To confirm this, we quantified ESRRB and SOX2 motif quality at each bound genomic site using FIMO ^59^ and compared them to the corresponding ChIP enrichment values (Figure 4F and Methods). For ESRRBwt, ChIP enrichment correlated negatively with SOX2 motif strength and showed only a weak positive association with the ESRRB motif. Conversely, ESRRBhydro displayed a positive correlation with SOX2 motif quality and a negative one with the ESRRB motif, indicating that ESRRBhydro preferentially associates with genomic regions containing high-quality SOX2 motifs. In contrast, SOX2wt bound its targets largely independently of motif quality, whereas SOX2hydro and SOX2neg showed a modest positive correlation with motif strength. SOX2neg binding also correlated positively, albeit weakly, with high-quality ESRRB motifs (Figure S6F). Analysis of endogenous SOX2 occupancy revealed higher SOX2 ChIP signal at ESRRBhydro rather than ESRRBwt regions (Figure 4G and Figure S6G). Taken together, these results show that the less search-efficient ESRRBhydro is redirected to SOX2 regions. As ESRRB and SOX2 can physically interact independently of DNA ^33^, our experiments suggest that SOX2 contributes to ESRRB’s target search and might guide ESRRB to defined genomic regions.

To further characterize the regions bound by vTFs, we extended the motif enrichment analysis to other TFs of the pluripotency network. The higher frequency of OCT4, OCT4::SOX2 and ESRRB motifs at SOX2hydro and SOX2neg peaks (Figure S6H) suggests that, because their search efficiency is reduced, they may preferentially associate with regions already occupied by other transcription factors. The OCT4::SOX2 and OCT4 motifs were also enriched at ESRRBhydro peaks, although to a lesser extent than the SOX2 motif. On the other hand, the motif for NR5A2, which often binds together with ESRRB ^30^ was frequently found at ESRRBwt peaks (Figure S6H). Integration of ChIP-seq and ATAC-seq data in G1 phase ^29^ (Figure S6I) further revealed that SOX2neg is bound to more accessible regions, which may suggest a loss of pioneering activity. Conversely, ESRRBhydro-bound sites are less accessible than ESRRBwt ones, in agreement with a model where SOX2 predominantly guides ESRRBhydro binding and target search.

Finally, we examined whether the vTFs retained the ability to sample unbound motifs by determining ChIP-seq enrichment at genomic regions containing SOX2 or ESRRB motifs but that were not bound or bound below the peak-calling threshold (^60^ and Methods). SOX2neg displayed a markedly reduced signal at unbound SOX2 motifs compared to SOX2wt and SOX2hydro (Figure 4H), suggesting a loss of ability to transiently sample cognate sites, consistently with its reduced number of binding trials and increase in diffusion time observed in single-molecule imaging (Figure 3H-I). This suggests that the short binding events of SOX2neg are depleted in actual binding to SOX2 motifs. These results are also in agreement with previously published results from our laboratory ^19^ and on a FOXA1 mutant with reduced positive charge ^60^. SOX2hydro sampling was not affected. On the contrary, ESRRBhydro showed reduced but above-background enrichment at unbound ESRRB sites compared to ESRRBwt, confirming that ESRRBhydro can still recognize its targets but then fails to maintain stable binding.

### SOX2 loss decreases ESRRB search efficiency and genome occupancy

To better understand the role of SOX2 in ESRRB search dynamics, we took advantage of the 2TS22C mES cell line ^61^ in which a TET-off system allows inducible SOX2 depletion upon dox treatment. After 26h of dox treatment (SOFF) that results in near-full loss of SOX2 ^62^ (Figure S7A-B), both ESRRB genomic enrichment and the number of bound regions were diminished (Figure S7C-E). Interestingly, ESRRB average ChIP-Seq enrichment score was reduced by 34% at all sites and slightly more (40%) at regions normally co-bound by SOX2 (Figure S7F), indicating that a portion of ESRRB binding is SOX2-dependent, and suggesting that SOX2 may facilitate efficient ESRRB chromatin association. ESRRB protein levels, instead, were only 12% lower than in control conditions (SON) (Figure S7B), indicating that changes in ESRRB binding upon SOX2 removal are not simply a consequence of lower ESRRB expression. Note that while secondary chromatin and pluripotency network changes may contribute to the observed ESRRB search phenotypes, we think this is unlikely since changes in ESRRB levels and other pluripotency TFs such as OCT4 are minimal in these conditions (^62^ and Figure S7B).

We next asked how search dynamics of ESRRBwt and ESRRBhydro are impacted by SOX2 loss. Because doxycycline is used to deplete SOX2, ESRRBwt and ESRRBhydro were expressed under an EF1α constitutive promoter, and labelled very sparsely for single-molecule imaging experiments. Comparing ESRRBwt and ESRRBhydro in the presence and absence of SOX2 allowed us to separate the contributions of their intrinsic search ability from cooperativity with SOX2 in their ability to locate their targets. Diffusion analysis (Figure S8A-B and Supplementary Tables 2 and 3) revealed that the total bound fractions of both ESRRBwt and ESRRBhydro were reduced upon SOX2 loss, while the fast diffusing population increased at the expense of the slow diffusing one (Figure 5A). Additionally, we registered increments in diffusion speed for all fractions (Figure 5B) when SOX2 is removed. Overall, this translates in an increase in D_eff_ of 2-fold for ESRRBwt and of 3-fold for ESRRBhydro, suggesting that SOX2 depletion increases ESRRB overall mobility (Figure S9A). Next, we questioned whether SOX2 also impacts ESRRB motion directionality. The angle distribution probability showed a small decrease in directional bias in the absence of SOX2 for ESRRBwt but not for ESRRBhydro (Figure S9B). Accordingly, the *f_180/0_* coefficients calculated from free diffusing tracks of ESRRBwt and ESRRBhydro were unchanged upon SOX2 depletion (Figure 5C), albeit we observed increased anisotropy at short displacements for both ESRRBwt and ESRRBhydro in absence of SOX2 (Figure S9C). Taken together, our results indicate that SOX2 slows down ESRRB nuclear diffusion, further increasing its presence in regions of higher chromatin density ^11^ and facilitating ESRRB interactions with DNA. On the other hand, the anisotropy of ESRRB diffusion is only marginally impacted by SOX2, and is mostly determined by its own biophysical properties. While these results are in line with SOX2 assisting ESRRB to find some of its target sites, we cannot exclude that ESRRBhydro redirection may partially reflect enhanced SOX2 tethering because of its increased hydrophobicity.

**Figure 5.**
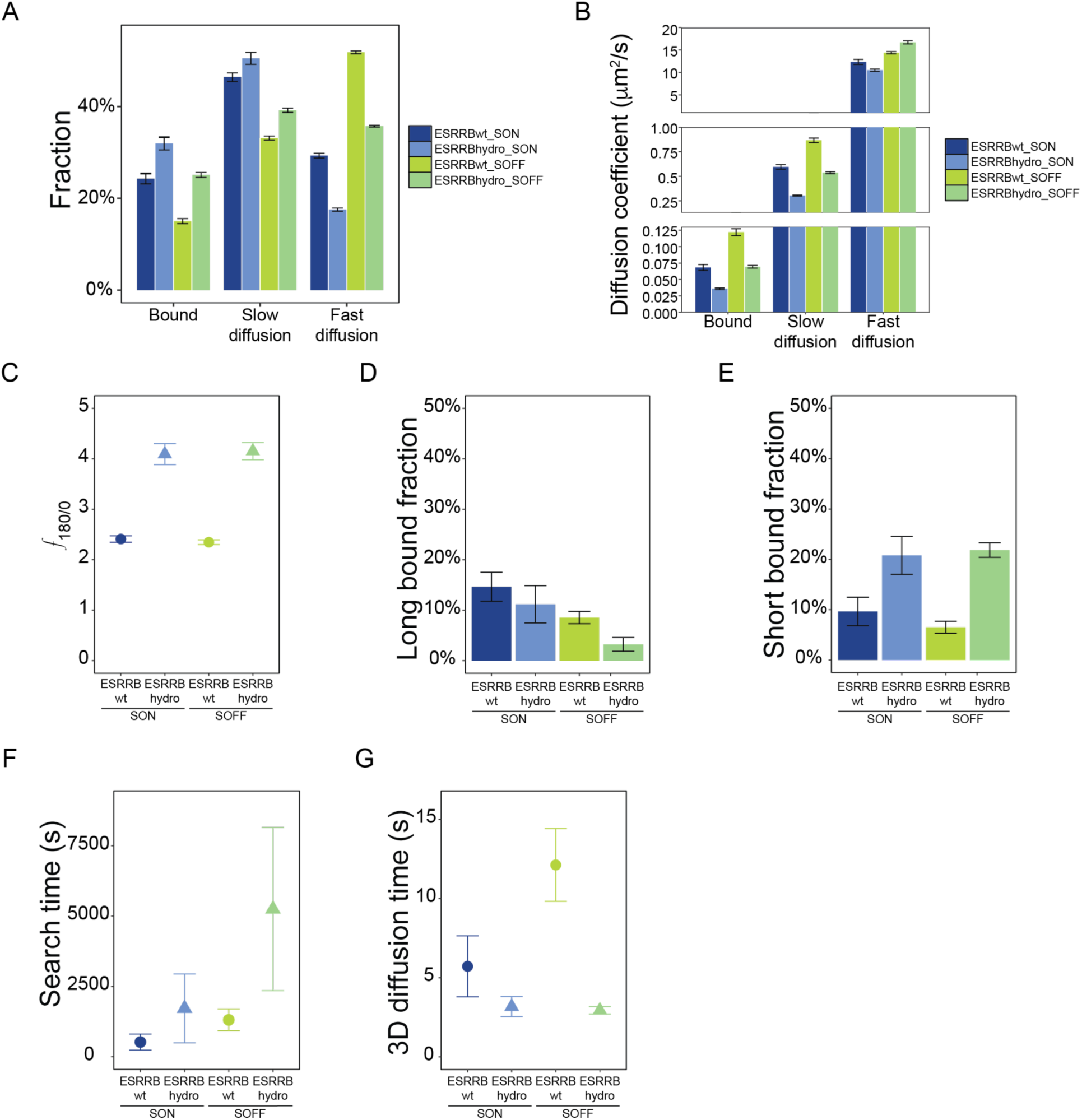
SOX2 supports efficient ESRRB search. **A**) Fractions of bound, slow and fast diffusing molecules, derived from fitting the continuous movies with a three-states diffusion model. Data are provided as mean values ± standard deviation of 500 resamplings with 80% of data. **B**) Diffusion coefficients of bound, slow and fast diffusing molecules. Data are provided as mean values ± standard deviation of 500 resamplings with 80% of data. **C**) Fold anisotropy ratio *f_180/0_*. Data are presented as mean values ± standard deviation. **D-E**) Average fractions of all binding events lasting less than 30 s (**D**) and more than 30 s (**E**). Data are provided as mean values ± standard deviation. **F**) Search time needed for a single TF molecule to find a specific site (Methods and Supplementary Table 7). **G**) Time spent diffusing between binding events (Methods and Supplementary Table 7). For panels G and H, data are provided as mean values ± error from error propagation.

To determine whether SOX2 primarily impacts ESRRB stable or rather transient interactions with DNA, we first plotted the cumulative binding curve (Figure S9D), then calculated the fraction of long (above 30 s) and short (below 30 s) binding events (Figures 5D-E) and the relative average residence times (Figures S9E-F). For both ESRRBwt and ESRRBhydro, SOX2 depletion led to a faster decline in the bound fraction across time, indicating fewer molecules capable of maintaining prolonged binding. This effect was more pronounced for ESRRBhydro (Figure S9D). Loss of SOX2 caused a greater reduction in the fraction of long binding events compared to an increase in hydrophobicity (Figure 5D). In contrast, the fraction of short binding events was unaffected by SOX2 loss (Figure 5E). In summary, our results indicate that SOX2 is dispensable for ESRRB’s non-specific DNA interactions, but that cooperativity with SOX2 may increase the fraction of long-binding events to ESRRB target regions.

Finally, we quantified the target search parameters (Figure 5F-G and Figure S9G). Upon SOX2 depletion, the overall search time increased for both ESRRBwt and ESRRBhydro, with a larger effect observed for the hydrophobic variant (Figure 5F). In the case of ESRRBwt, the increase was primarily driven by a longer 3D diffusion time between binding trials (Figure 5G), indicating that in the absence of SOX2, ESRRB spends more time freely diffusing before locating target sites. In contrast, the higher search time of ESRRBhydro was mainly due to an increased number of binding trials (Figure S9G), highlighting how high hydrophobicity, in absence of an interacting TF, hinders the efficiency of TF search.

### Cooperative binding of SOX2 and ESRRB at shared target regions

We have shown that ESRRB search is enhanced by the presence and the interaction with SOX2. We next examined how SOX2 affects ESRRB genomic binding, and conversely, whether ESRRB influences SOX2 binding despite its minimal impact on SOX2 search dynamics. We made use of our data in 2TS22C (Figure S7D) and previously published data in an ESRRB inducible mESCs model (EKOiE cells ^30^) to identify regions losing ESRRB or SOX2 binding upon SOX2 or ESRRB depletion, respectively. We first classified ESRRB-bound regions according to changes in ESRRB occupancy in the presence and absence of SOX2 in 2TS22C cells. We used a log2 fold-change cutoff of ± 0.5 (about 40% difference between conditions) to distinguish regions losing (cluster E1) or gaining (cluster E2) ESRRB binding upon SOX2 depletion, considering all leftover regions as unchanged (cluster E3) (Figure 6A). In absence of SOX2, ESRRB binding was reduced or lost at 21942 target regions, while 13857 were unaffected. Only a small subset of regions (925) increased ESRRB occupancy. We then carried out the same analysis on SOX2 targets upon dox-induced ESRRB removal (EOFF) and in control conditions (EON). We identified regions losing or reducing SOX2 binding upon ESRRB depletion (cluster S1, 26703 regions), regions gaining SOX2 binding (cluster S2, 7932 regions) and SOX2 loci not affected by ESRRB loss (cluster S3, 37568 regions). Overall, ESRRB binding was lost at about 60% of its target loci upon SOX2 depletion, while half of SOX2 sites were not affected by ESRRB removal, and only 38% lost SOX2 binding (Figure 6C). This suggests an asymmetric relationship where ESRRB binding depends more on SOX2 than vice-versa. The genomic feature distribution was broadly similar across clusters, with only a modest bias towards enhancer-associated regions in clusters E2 and S2 and promoter-associated regions in cluster E3 and S3 compared to the other clusters (Figure S10A). As expected, clusters E1 and S1 had a slightly higher frequency of SOX2 and ESRRB motifs respectively (Figure S10B-C).

**Figure 6.**
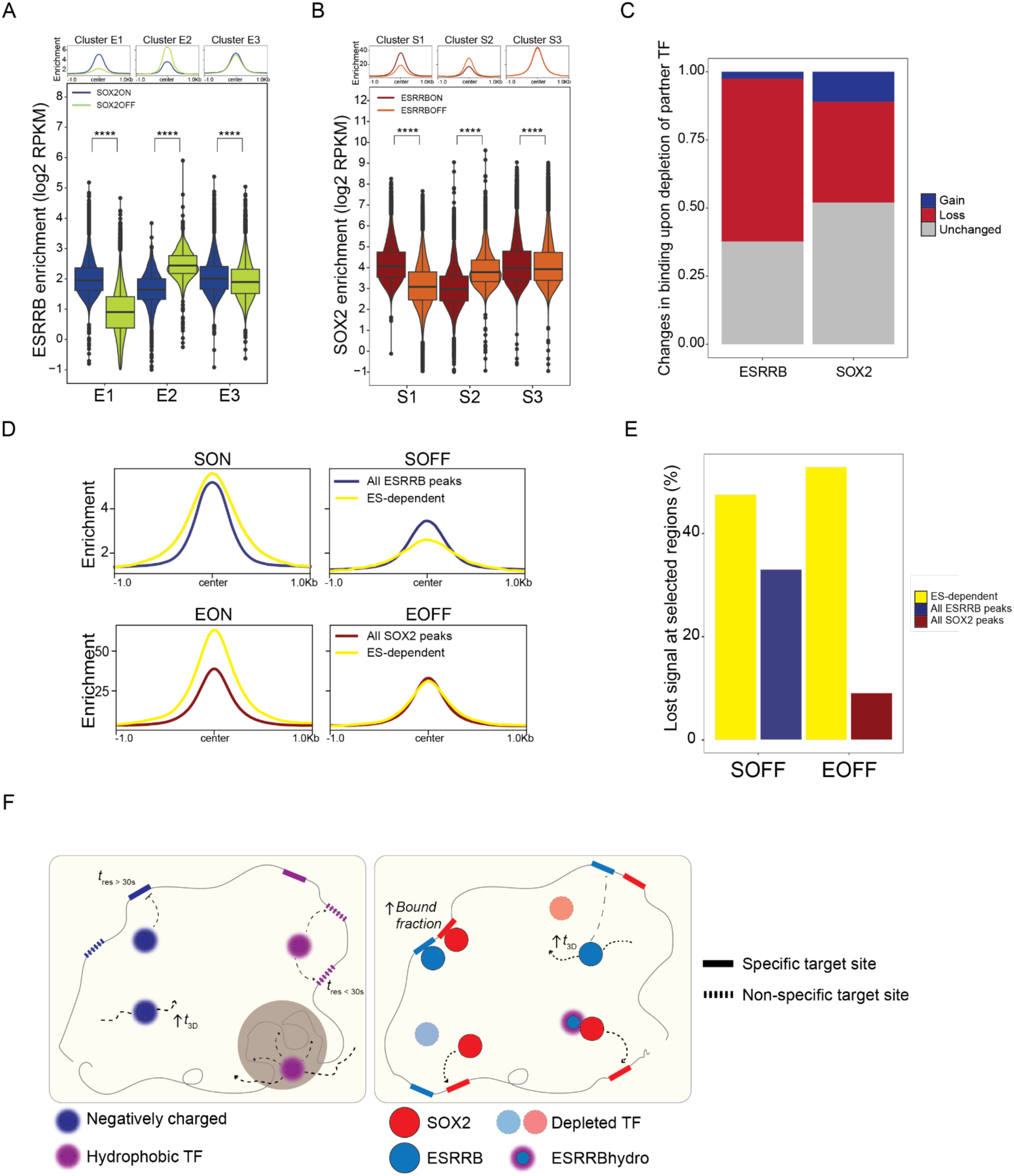
ESRRB and SOX2 mutually stabilize their binding at a subset of pluripotency regulatory elements. **A**) Average enrichment score (log2 RPKM) of ESRRB ChIP-Seq at cluster regions in SON and SOFF conditions. **B**) Average enrichment score (log2 RPKM) of SOX2 ChIP-Seq at cluster regions in EON and EOFF conditions. Data from ^30^. **C**) Fraction of ESRRB sites (ESRRB) and SOX2 sites (SOX2) sites gaining, losing or not changing binding upon SOX2 or ESRRB loss respectively. **D**) ESRRB enrichment profiles in SON and SOFF conditions (top) and SOX2 (bottom) in EON and EOFF conditions. Comparison of enrichment profiles at ES-dependent regions (yellow) against global ESRRB and SOX2 enrichment profiles (see Figure S10D). **E**) Percentage of ESRRB signal lost upon SOX2 depletion (SOFF) and of SOX2 signal lost upon ESRRB depletion (EOFF) at ES-dependent regions (yellow) compared to global signal lost at all ESRRB peaks (blue) or at all SOX2 peaks (red) in SOFF and EOFF conditions respectively. **F**) Global scheme.

To gain further insights on SOX2 and ESRRB direct cooperativity, we crossed E1 and S1 clusters and identified 5199 regions where both TFs lose binding when the other is lost. These regions have high SOX2 and ESRRB binding in control conditions (Figure S10D), but lose 48% of ESRRB and 53% of SOX2 binding upon SOX2 and ESRRB depletion respectively, which is substantially more than the average loss of binding of 33% at all ESRRB sites and 9% at all SOX2 sites (Figure 6D-E and Figure S10D). Interestingly, binding enrichment of both SOX2 and ESRRB at these ES-dependent regions was higher than the genome average (Figure 6D), despite the quality of the SOX2 and ESRRB motifs at ES-dependent sites being slightly poorer than the the average global ESRRB motif quality (Figure S10E and Methods). Overall, this suggests that ESRRB and SOX2 cooperatively reinforce each other’s binding at a subset of regulatory elements with weaker motifs.

## Discussion

This study investigates the relative contribution of intrinsic biophysical properties and cooperative interactions to TF search and binding. We focused on ESRRB and SOX2, both highly expressed in mESCs and fundamental for the sustenance of the naïve pluripotency network ^28^. By altering TF hydrophobicity and electrostatic charges and comparing microscopy and genomics data with wild-type proteins, we were able to dissect the impact of these protein features on search parameters. Additionally, we assessed the importance of TF cooperativity by comparing SOX2 and ESRRB genomic occupancy in wild-type conditions and upon depletion of the other TF, and ESRRB search in presence and absence of SOX2. Our results reveal that, while sharing a good subset of target regions ^30,63–66^, SOX2 and ESRRB employ strikingly different search strategies, highlighting the TF-specific nature of the search process.

The novelty of our approach consists in modulating hydrophobicity or net charge by adding short amino-acid stretches to the C or N termini of our TFs, without alteration of the native coding sequence itself. This strategy, contrary to point mutations ^67,68^ and domain swapping ^13,69^, has the advantage of minimizing the bias of introducing or replacing protein interaction modules. Although we cannot fully exclude minor effects on cofactor binding, this design reduces the likelihood of inadvertently recruiting different cofactors or creating novel interactions. The addition of hydrophobic or negatively charged amino acidic stretches may push ESRRB and SOX2 biophysical properties at the limit or even beyond their physiological range, as in the case of ESRRBhydro. Nevertheless, such perturbations were instrumental to isolate and interrogate the specific contribution of individual parameters to TF functionality. By amplifying these features, we were able to resolve how distinct biophysical traits shape the overall search and binding behavior of ESRRB and SOX2. We show that SOX2 and ESRRB exhibit fundamentally different search strategies, with SOX2 maintaining robust search efficiency even upon perturbation of its electrostaticity/hydrophilicity. ESRRB, in contrast, is highly sensitive to increased hydrophobicity and to SOX2 loss, which may indicate a less robust, more cooperativity-dependent search strategy.

SOX2 is perhaps the most extensively studied TF in the single-molecule imaging field ^12,19,32,33,45,60^. Compared to early works by ^32,33,45,60^ where residence times were extracted by fitting single particle trajectories with a double-exponential decay equation, we chose to use GRID ^19,44,51^. This approach has the advantage of resolving distinct binding behaviours in an unbiased manner. We fixed an arbitrary threshold at 30 s, and derived the corresponding average long and short residence times by averaging values above and below the threshold. This led to an average long residence time for SOX2wt (260 s) which is in line with a previous study from our laboratory ^19^. A similarly average long residence time was reported for SOX2 by live imaging in the mouse embryo ^70^. In recent years hydrophobicity is emerging as a key determinant of protein-DNA interactions, bridging unspecific electrostatic interactions and specific amino acids - nucleotides hydrogen bonds as TFs strive to bind to their motifs ^71,72^. Consistent with the fundamental laws of physics, we show that hydrophobicity overall slows down TF mobility in the nucleoplasm and facilitates confinement. This difference is more evident for SOX2 than ESRRB, possibly because ESRRBwt is already approximately three times more hydrophobic than SOX2 (Figure 1c). The observation that around 20% of SOX2hydro, ESRRBwt and ESRRBhydro molecules are engaged in short, transient binding events (< 30s) might reflect both the increased chances of protein-protein and protein-DNA contacts in chromatin domains ^11^ and the role of hydrophobicity in facilitating TF-TF interactions ^73–75^. Overall, the effect of hydrophobicity on target search efficiency appears to be modulating the balance between specific and nonspecific interactions. As we show in particular with ESRRBhydro, excessive hydrophobicity disrupts this balance. By shifting the proportion of short and long interactions, hydrophobicity becomes detrimental by causing TFs to become unproductively trapped, as evidenced by the discrepancy between long bound fraction and ChIP-Seq peaks for SOX2hydro and ESRRBhydro.

In addition to hydrophobicity, we also investigated the impact of electrostatic charges on search dynamics with the SOX2neg variant. SOX2 carries a net positive charge and is about three times more positively charged than ESRRB. We show that changes in electrostatic properties do not impact SOX2 anisotropy but do cause a reduction of the long bound fraction and corresponding residence time, together with an increase of the time spent diffusing between binding events (t3D). The reduced search efficiency of SOX2neg correlates with markedly lower genomic occupancy and an impaired ability to sample unbound sites and bind motifs located in low accessibility regions. This is in agreement with the current knowledge on the role of electrostatic interactions in TF target search ^19,76,77^ and with very recent findings that phosphorylation of SOX2 during mitosis reduces its pioneering capacity (Williams et al., 2025). Regions bound by SOX2neg are also more enriched for TF motifs (ESRRB, NR5A2, NANOG and OCT4::SOX2), suggesting that cooperative binding or higher motif density might compensate for reduced search efficiency In the second part of this paper, we focus on the relationship between ESRRB and SOX2 and the contribution of SOX2 to ESRRB’s target search. These two TFs belong to different families ^1^, are both important pluripotency factors but also have distinct roles and partners in development ^78,79^. They can interact through their DNA-binding domains even in the absence of DNA ^33^. Here we build on previous work on ESRRB and SOX2 search dynamics in presence and absence of their partner ^33^ and determine how their interaction during the search process translates in binding occupancy. Leveraging on our ESRRBhydro mutant whose search and binding capacities are compromised, and comparing ESRRBwt and ESRRBhydro dynamics in the presence or absence of SOX2, we were able to disentangle which aspects of ESRRB’s search rely on SOX2 and which are determined by ESRRB’s intrinsic biophysical properties. Upon SOX2 depletion, ESRRB becomes more mobile and spends a larger fraction of time diffusing between binding attempts. The short-bound fraction of ESRRB is primarily determined by its hydrophobicity, whereas the long-bound fraction is more impacted by the presence of SOX2. Overall, these observations indicate that SOX2 plays a major role in facilitating ESRRB search. Accordingly, ESRRB genomic occupancy is strongly reduced in the absence of SOX2. SOX2 binding, instead, is only partially affected by the loss of ESRRB, mainly at a subset of pluripotency-related regulatory regions. This suggests that, while ESRRB does not play a role in SOX2 search, it supports its binding to lower affinity sites in regions where they bind together. The dominant role played by SOX2 is consistent with its ability to redirect TFs to different sets of regulatory regions ^29^. Our study represents an important step in our still limited understanding of how hierarchical and cooperative TF interactions modulate target search.

## Material and methods

### Cell culture

CGR8 and 2TS22C ^13^ mouse embryonic stem cells were routinely cultured on 0.1% gelatine (Sigma, G9391) - coated dishes at 37 °C and 5% CO_2_. CGR8 and 2TS22C were grown in GMEM (Sigma, G5154) supplemented with 10% heat-inactivated fetal bovine serum (Gibco, 16141-079), 1% nonessential amino acids (Gibco, 11140–050), 2 mM L-glutamine (Gibco, 25030–024), 2 mM sodium pyruvate (Sigma, S7636), 100µM 2-mercaptoethanol (Sigma, M3148), 1% penicillin and streptomycin (BioConcept, 4–01F00-H), in-house produced leukemia inhibitory factor, 3 µM CHIR99021 (Sigma, 361559) and 0.8 µM PD184352 (Sigma, PZ0181).

Cells were split every 2-3 days by trypsinization (Sigma T4049). To induce vTF expression cells were treated with Doxycycline (Sigma, D3447) 24h before experiments at the following concentrations: 500 ng/mL for FACS, flow cytometry and immunofluorescence, 10-50 ng/mL for single molecule imaging experiments. SOX2 was depleted in 2TS22C cells by treating them with 1μg/mL of doxycycline for 26 h. For imaging experiments, cells were plated the day before on laminin – coated dishes (BioLamina, LN511-0202) on a 1:10 dilution in PBS. Prior to imaging, GMEM was replaced with Fluorobrite DMEM (Thermo Fisher Scientific, A18967-01), 2 mM sodium pyruvate, 1% non-essential amino acids, 1% penicillin and streptomycin (BioConcept, 4-01F00H), 2 mM L-Glutamine and 100 µM 2-mercaptoethanol, supplemented with FBS, 3 µM CHIR99021 and 0.8 µM PD184352 for CGR8 and 2TS22C.

### Plasmid construction

cDNA sequences for ESRRB, SOX2 and Halo-Tag were amplified from the pLV-TRE3G-ESRRB-SNAP ^80^ and the pLV-TRE3G-Halo-SOX2 ^81^ plasmids present in the lab. The HA-Tag sequence was integrated into the constructs by including it into one of the primers (Table S1). The resulting ESRRB, SOX2-HA and Halo-HA fragments were cloned into the pLV-TRE3G-MCS plasmid ^81^ to generate the pLV-TRE3G-ESRRBwt-Halo-HA and pLV-TRE3G-Halo-SOX2wt-HA plasmids. Sense and reverse complements of single-stranded DNA oligonucleotides coding for hydrophobic or negatively charged amino acids were annealed, purified and cloned using the ClaI (NEB, R0197S) restriction enzyme to obtain the pLV-TRE3G-ESRRBhydro-Halo-HA, pLV-TRE3G-Halo-SOX2hydro-HA and TRE3G-Halo-SOX2neg-HA plasmids. To generate the pLV-EF1a-ESRRBwt-Halo-HA and pLV-EF1a-ESRRBhydro-HA plasmids, the entire ESRRBwt-Halo-HA and ESRRBhydro-Halo-HA sequences were amplified and introduced into the pLV-EF1a-mCherry plasmid via In-fusion cloning (Takara Bio, 638947) while removing mCherry. All constructs were sequence-verified by Sanger Sequencing (Microsynth). Primer sequences used in this study are available in Table S1.

### Lentiviral vector production

Lentiviral vectors were generated by calcium-phosphate transfection ^82^ of HEK 293T cells with the envelope plasmid psPAX2 (Addgene, 12260), the packaging plasmid pMD2.G (Addgene, 12259) packaging plasmid ^83^ and the lentiviral construct of interest. Cell media were collected two days after transfection and lentiviral vectors were concentrated by ultracentrifugation for 2 h at 20′000 g and 4°C. Aliquots of 50 µL were immediately used for transduction or stored at −80 °C.

### Generation of Halo cell lines

CGR8 cells previously engineered to express the rtTA3G ^19^ were transduced with 50 µL of concentrated lentiviral particles carrying one of the TF-Halo-HA constructs and selected with 2 µg/mL of Puromycin (Gibco, A11138-03) for 7 days. To generate monoclonal cell lines, transduced cells were treated with 500 ng/mL Doxycline for 24h and stained with 100 nM Halo-TMR dye (Promega, G8252) for 30 minutes prior to sorting of single TMR-positive cells. Clones used in this study were selected by immunofluorescence based on Halo-TMR expression. EKO cells were first transduced with lentiviral particles containing the pLV-PGK-rtTA3G construct and selected with 5 µg/mL Blasticidin for one week, and were subsequently transduced with the TRE3G-SOX2wt-Halo-HA, TRE3G-SOX2hydro-Halo-HA and TRE3G-SOX2neg-Halo-HA lenviral vectors and selected for one week with 2 µg/mL puromycin. 2TS22C cells were transduced with lentiviral particles containing the EF1a-ESRRBwt-Halo-HA and EF1a-ESRRBhydro-HA constructs and selected with 2 µg/mL puromycin for one week. The resulting polyclonal cell lines were used for single molecule imaging experiments and constantly maintained under selection with puromycin.

### Immunofluorescence staining

Cells were plated on laminin (BioLamina, LN511-0202) diluted 1:10 in PBS +/+ (Gibco, 14040091) in 96-well plates (Pierce), at a density of 4000 cells /well. The next day, cells were fixed with 2% formaldehyde (Thermo Fisher Scientific, 28908) in PBS for 30 min at RT, permeabilized with 0.5% Triton X-100 (AppliChem, A1388,0500) in PBS for 30 min at RT then blocked with 1% BSA (Sigma, A7906) in PSBS. Cells were then incubated overnight at 4°C with one of the following primary antibodies diluted in 1% BSA-PBS: 2 µg/mL anti-ESRRB (BioTechne, PP-H6705-00), 1:200 anti-NANOG (Abcam, ab80892), 1:500 anti-OCT4 (Cell Signaling Technology, 5677), 1:1000 anti-HA.11 (BioLegend, 901501). The following morning, cells were washed twice with PBS, before addition of the secondary antibodies Alexa-Fluor-647 goat anti-rabbit IgG H+L (Thermo Fisher Scientific, A21244) or Alexa-Fluor-647 Donkey anti-mouse IgG H+L (Thermo Fisher Scientific, A31571) diluted 1:1000 in 1% BSA-PBS for 1h at RT. Cells were then washed twice with a solution of 0.1% Tween-20 (Thermo Fisher Scientific, 10113103) in PBS and twice with PBS before addition of DAPI-Fluoromount G (SouthernBiotech, 0100-20). For quantification of TF levels, imaging was performed on an INCell Analyser 2200 (GE Healthcare) with a 20X magnification objective, 30 ms exposure for DAPI channel and 50ms for the Cy5 channel.

### ChIP-Seq

All experiments were performed in duplicates, with the exception of the SOX2 and ESRRB ChIP in 2TS22C SON that were performed in triplicates and the endogenous ESRRB and SOX2 ChIP in the ESRRBwt-Halo and the SOX2wt-Halo cell lines for which only one replicate was performed. For all experiments, cells were collected after trypsinization, centrifuged, resuspended in PBS and incubated with 2mM DSG (Thermo Fisher Scientific, 20593) for 50 min at RT on a rotating platform. After centrifugation, a second fixation was carried out with 1% formaldehyde for 10 min at RT. The reaction was quenched with 200 nM Tris-HCl pH 8.0 (PanReac AppliChem, A4577) for 10 min. To sort CGR8 cells with the same TF-Halo mean fluorescence intensity, trypsinized cells were stained with 100 nM Halo-TMR and 40.5 µM Hoechst 33342 (Thermo Fisher Scientific, H3570) diluted in GMEM medium for 30 minutes on a rotating platform at room temperature (RT) prior to fixation. For nuclear extraction, cell pellets were incubated twice with LB1 (50 mM HEPES-KOH pH 7.4, 140 mM NaCl, 1mM EDTA, 0.5 mM EGTA, 10% Glycerol, 0.5% NP-40, 0.25% Triton X-100) supplemented with a 1:100 dilution of Protease Inhibitor Cocktail (Sigma, P8340-1ML), incubated for 10 min at 4°C with 100 rpm gentle shaking and spun down for 5 min at 4°C and 1,700 g. Resulting nuclei were then resuspended in LB2 (10 mM Tris-HCl pH 8.0, 200 mM NaCl, 1 mM EDTA, 0.5 mM EGTA, 1:100 Protease Inhibitor Cocktail), incubated for 10 min at 4°C with 100 rpm gentle shaking, and spun down for 5 min at 4°C and 1700 g. Nuclei were washed twice with SDS shearing buffer (10 mM Tris-HCl pH 8.0, 1 mM EDTA, 0.15% SDS, 1:100 Protease Inhibitor Cocktail) without disturbing the pellet and finally resuspended in SDS shearing buffer. Chromatin was immediately sonicated for 20 min at 5% duty, 140 W, 200 cycles, in a Covaris E220 focused ultrasonicator. Chromatin was then centrifuged for 5 min at 4°C and 10,000 g to remove the insoluble material, and the supernatant was transferred to a new tube. As total input control, 18 μL of chromatin were incubated first with 1X TE buffer pH 8.0 (PanReac AppliChem, A0386) and 10 ng/μL RNAse A (Qiagen, 19101) for 30 min at 37°C, then with 400 ng/μL of Proteinase K (Qiagen, 19131) for 30 min at 55°C and 1,100 rpm shaking. To reverse the crosslinking, 200 mM NaCl (Sigma, 59222C) was added and samples were incubated at 65°C for 16 h and 1,100 rpm shaking. Chromatin was purified using a MinElute PCR purification kit (Qiagen, 28004). ChIPs were performed on 3.5 μg of chromatin using the ChIP-IT High Sensitivity kit (Active motif, 53040) according to manufacturer’s instruction with 5 μg of anti-HA.11 (Biolegend, 901501) or 5.8 μg of anti-ESRRB (BioTechne, PP-H6705-00) or 3.1 μg of anti-SOX2 (Cell Signalling Technology, 23064) antibodies. 20 ng of Drosophila spike-in chromatin (Active motif, 53083) and 2 μg of spike-in antibody (Active motif, 61686) were added for internal normalization ^84^. Sequencing libraries were prepared using the NEBNext Ultra II DNA Library Prep Kit (NEB, E7645L) and sequenced on an Illumina NextSeq 500 sequencer using 75-nucleotide read length paired-end sequencing or on a Element Biosciences Aviti sequencer.

### Single-molecule imaging set-up

Single-molecule imaging experiments were performed using a Inverted Nikon Eclipse Ti2-E motorized microscope equipped with a 100x/1.49 oil immersion SR HP APO-TIRF objective and a Photometrix 95B sCMOS camera to detect fluorescent emissions. Imaging was performed with a HILO (highly inclined and laminated optical sheet) ^85^ set up using a 561nm diode laser for Halo-JF549 excitation, and 405nm diode laser for Hoechst excitation, with a 405/488/568/647 DM wheel emission filter. For slow SMT, in order to correctly differentiate between unbinding of molecules and photobleaching we recorded 5 min - long movies keeping a constant exposure time of 60 ms and total time intervals (ti) of 150 ms, 240 ms, 420 ms, 600 ms and 5 s. Laser power was set at 13mW. To study diffusion coefficients of fast-moving molecules, we recorded 1 min - long movies with a 10ms exposure time at 110mW laser power. Data were acquired in two to three biological replicates. Statistics are reported in Supplementary Tables 2-7.

### In silico protein analysis

The R package Peptide ^13^ was used to compute TF physicochemical properties. Specifically, the net charge was calculated using the function charge with parameters pH = 7.2 and pKscale = “EMBOSS”. The GRAVY (Grand Average of Hydropathy) index was computed with the function hydrophobicity and setting scale = “KyteDoolittle”. The Kyte and Doolittle scale assigns each amino acid a numerical hydrophobicity value based on experimental observations of how it behaves in water versus non-polar solvents ^43^. Amino acid sequences for the SOX and ERR mouse TF families were collected using the R package UniProt.ws 2.50.0 ^14^ with taxId = 10090. 3D protein structures and interactions with DNA were modelled with Alphafold 3 ^42^ using ColabFold ^86^.

### Immunofluorescence analysis

Images were processed using the CellProfiler software ^87^. Metadata were extracted from the image files recovering the following information: well, field, imaging channel, cell line and target protein. Processing was performed on the DAPI channel image to identify nuclei and create segmentation masks. Images were first rescaled to use the full intensity range, followed by median filtering (artifact diameter =5) to enhance contrast and reduce imaging artifacts. Primary objects (nuclei) were identified and filtered to be between 20 and 35 pixels in diameter. Further filtering was done selecting minimum cross-entropy thresholding with 1.3488 smoothing scale, 2.4 correction factor and 0.1 and 1 for lower- and upper-bound thresholds, respectively. Cell clumps were identified using the shape function and removed from the analysis. uclei were further filtered by eccentricity (0-0.8) and compactness (above 1.3) to exclude irregular shapes. The resulting mask was used to measure integrated intensity within each nucleus across all imaging channels.

### ChIP-Seq analysis

ChIP-Seq libraries (newly generated or published data) and published ATAC-Seq ^62^ libraries were mapped to the mm10 version of the mouse genome and to the BDGP6 version of the Drosophila melanogaster genome with STAR 2.7.11 ^88^ setting parameters --runMode alignReads --alignMatesGapMax 2000 --alignIntronMax 1 --alignEndsType EndToEnd --outFilterMultimapNmax 1. Duplicate reads were removed using Picard (Broad Institute), then samples were indexed using SAMTools 1.19.2 ^89^. Deduplicated bam files were spike-in normalized based on the sample with the lowest number of Drosophila reads ^90,91^ and downsampled using SAMTools. Biological replicates were merged using SAMTools, and peaks were called on merged bam files using MACS3 2.2.4 ^92^ with settings -f BAMPE -g mm and merged with awk. Blacklisted peaks ^93^ were removed from the final peak set using the BEDTools 2.29.2 ^94^ intersect command with option -v. Common peaks between samples were identified with bedtools intersect with option -wa.

Bigwig files were generated starting from replicate-merged bam files using the deepTools 3.5.1 ^95^ bamCoverage and the setting --normalizeUsingRPKM. To compare enrichment scores between samples, peak files were merged to generate a consensus bed file using BEDOPS 2.4.41 ^96^, and then enrichment scores were calculated in these regions using the multiBigwigSummary function of deepTools with the setting BED-file.

Heatmaps were generated using the deepTools function plotHeatmap. Enrichment profiles were generated with a custom R code starting from data tabs derived by the deepTools function plotProfile --outFileNameData.

### Published datasets

SOX2 ChIP-Seq data in presence/absence of ESRRB were obtained from GSE152186 ^30^ and SOX2 ChIP-Seq data in 2TS22C were obtained from GSE241731 ^29^. ATAC-Seq data in CGR8 cells were obtained from GSE241731 ^29^.

### Motif analysis

Motif analysis was performed with HOMER 5.1 ^58^ using the command findMotifsGenome.pl on replicate-merged peak sets with default parameters. Motif frequency was plotted for the following Homer motifs: 95 (ESRRB); 338 (SOX2); 256 (OCT4::SOX2); 254 (OCT4); 410 (NANOG); 247 (NR5A2). Motif quality was determined by analysing peaks files with FIMO ^59^ with settings –thresh 0.001 and JASPAR ^97^ motifs MA0141.1 (ESRRB) and MA0141.3 (SOX2). Assessment of SOX2 and ESRRB sampling was done as previously described in ^60^. Briefly, we scanned the mouse mm10 genome version for the ESRRB (Esrrb(NR)/mES-Esrrb-ChIP-Seq(GSE11431)/Homer - motif 95) or SOX2 (Sox2(HMG)/mES-Sox2-ChIP-Seq(GSE11431)/Homer - motif 338) position weight matrices to identify all occurrences of ESRRB or SOX2 motifs. Background sequences not overlapping ESRRB or SOX2 motifs were generated with the command bedtools shuffle -chrom. To identify unbound sites, the ESRRB or SOX2 motifs sets were crossed with the replicate-merged ESRRB (ESRRBwt + ESRRBhydro) or SOX2 (SOX2wt + SOX2hydro + SOX2neg) peaks sets with bedtools intersect -v. Enrichment over unbound motifs and background control sequences was calculated using merged bw files and the deeptools command computeMatrix reference-point --referencePoint center --missingDataAsZero as described in ^60^.

### Single molecule data analysis

Nuclei were detected in Fiji using the Stardist plugin ^98^, and single cell movies were cropped based on the nuclei ROI. The TrackIT ^44^ software was used to localize and track single molecules and to perform diffusion analysis. Spots were detected with a threshold of 0.7 for JF549 (CGR8 cells) and 0.6 for JF646 (2TS22C cells). For continuous movies, molecules were tracked through consecutive frames using the following parameters: tracking radius 8.1535 pixels, min. length before gap frame 2, min. track length 2, 1 gap framed allowed. For time-lapse imaging movies, the tracking radii were set as follows: 0.57 (150 ms ti), 0.8 (240 ms ti), 1.1 (420 ms ti), 2.3 (600 ms ti) and 3.37 (5 s ti), allowing respectively 3, 2, 2, 1 and 1 gap frames, with min. length before gap frame 2, min. track length 2. Tracking data statistics are available in Supplementary Table 2. Localization error was calculated at 0.0434 µm for JF549/CGR8 cells and at 0.0489 µm for JF646/2TS22C cells.

Diffusion rates and amplitudes were determined using TrackIT. Tracking data were fitted with a three-state brownian diffusion model ^31^, setting the bin width to 1nm. To avoid overestimation of bound molecules, only the first 10 jumps were retained for analysis, and jumps over gap frames were excluded. Errors of D_1,2,3,eff_, diffusion coefficients and A_1,2,3_ fractions were estimated by 500 repetitions of the diffusion analysis keeping only 80% of randomly chosen jump distances ^49^. Fraction A_1_ consists in the total (short + long binding events) bound fraction *f_b_*. The unbound fraction of tracked molecules can be calculated by *f_u_ = 1 - f_b_*. The average diffusion coefficient D_eff_ was determined as *∑_i_ D_i_* · *A_i_*. Diffusion statistics are available in Supplementary Table 3.

To study diffusional anisotropy, track segments were HMM-classified as bound or free using vbSPT ^50^, with default parameters and setting a minimum trajectory length of 3. To estimate the residual bound fraction in the free trajectories datasets, an automatic bound jump distance threshold was calculated assuming Brownian motion starting from the bound-state fraction (A1) and diffusion coefficient (D1) obtained from fitting the total population on TrackIT with the setting “Jumps to consider per track” = Inf. The residual fraction of bound molecules in the free pool was then computed by integrating the free subset probability distribution function (PDF) below this threshold. The jump angle anisotropy metric *f_180/0_* was calculated for each jump according to the formula 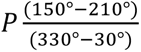 using TrackIt. To plot *f_180/0_* vs the mean displacement (Figure 2G and Figure S9C), the bin size was set to 25.

Binding kinetics and search time calculations were analysed and calculated as described in ^49,53^. Dissociation rate spectra were inferred from fluorescence survival time distributions obtained in time-lapse imaging experiments using GRID ^51^. This approach applies an inverse Laplace transformation to the survival time distributions, thereby resolving the amplitudes of dissociation rates *k_off,i_* associated with distinct binding states. Data from all time-lapse imaging conditions were pooled together for GRID analysis, and the spectrum of dissociation rates was obtained in the logarithmic range *log(k) = −3* to *log(k) = 1*. The error was estimated by repeating the GRID analysis 500 times with 80% of randomly selected survival time distributions. We defined long-binding events as having a maximal dissociation rate *k_off,min_* = 0.0333 s^-1^, corresponding to a minimum residence time of 30 s. The absolute fraction of long binding events *p_b,s_* was then calculated as 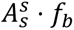, and the fraction of short binding events as 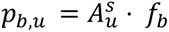, where 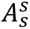 and 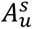 are the average amplitude of the GRID state spectrum ^51^ below and above *k_off,min_* ^49,53^. To quantify binding persistence, the bound fraction derived from diffusion analysis was weighted by the amplitudes of the GRID state spectra to account for the relative contribution of binding events with different lifetimes. The resulting weighted bound fraction was plotted as a function of residence time to generate the cumulative bound fraction, and represents the fraction of molecules that remain bound for at least the indicated residence time. Residence times were also obtained from the GRID state spectra. From the amplitude of the event spectrum below *k_off,min_* the number of non-specific binding events *N_trails_* can be calculated as 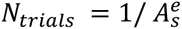. Target search time parameters were calculated assuming an equilibrium binding model with specific and unspecific binding sites as described in ^11,53^. Briefly, being the model described as:

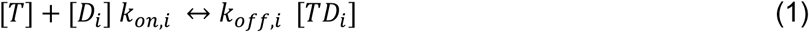

Where [*T*] is the concentration of free TF, [*D_i_*] the concentration of binding sites and [*TD_i_*] the concentration of TF-bound sites. At large concentrations of binding sites the pseudo association rate 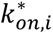 can be calculated as

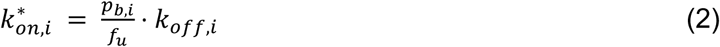

The target search time *t_search_* of a TF is determined by the number and the duration of nonspecific binding events before a specific target sequence is derived according to the relationship:

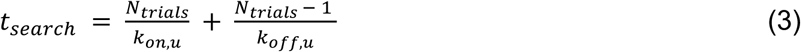

According to the facilitated diffusion model ^4^, the time a TF spends diffusing can be derived as

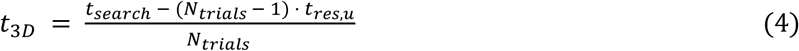

Where *t_res_*_,*u*_ is the residence time of unspecific binding events. Therefore

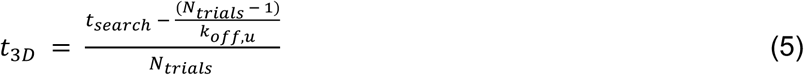

Time-lapse imaging tracking and search statistics are available in Supplementary Tables 3-4.

### Statistics

Results are presented as mean ± standard deviation unless otherwise stated.

## Acknowledgments

This study was founded by a Swiss National Science Foundation Sinergia grant (#CRSII5_189910) to D.M.S. and a post-doctoral fellowship to L.V. from the Peter und Traudl Engelhorn Stiftung. We thank the EPFL Gene Expression Core Facility for next-generation sequencing, the EPFL Flow Cytometry Core Facility, especially Francesco Palumbo and Gabriele De Simone, for cell sorting and the EPFL Bioimaging and Optics Platform, particularly Nicolas Chiaruttini.

## Authors Contributions

Conceptualization, LV and DMS; Methodology, LV and DMS; Formal Analysis, LV and DMS; Investigation, LV, CD and LF; Resources, DMS; Funding Acquisition, DMS; Supervision, DMS.

## Competing interests

The authors declare no competing interests.

## Notes

### Competing Interest Statement

The authors have declared no competing interest.

