## Supplementary Figures and Tables for "The role of charge, hydrophobicity, and cooperativity in target search of SOX2 and ESRRB"

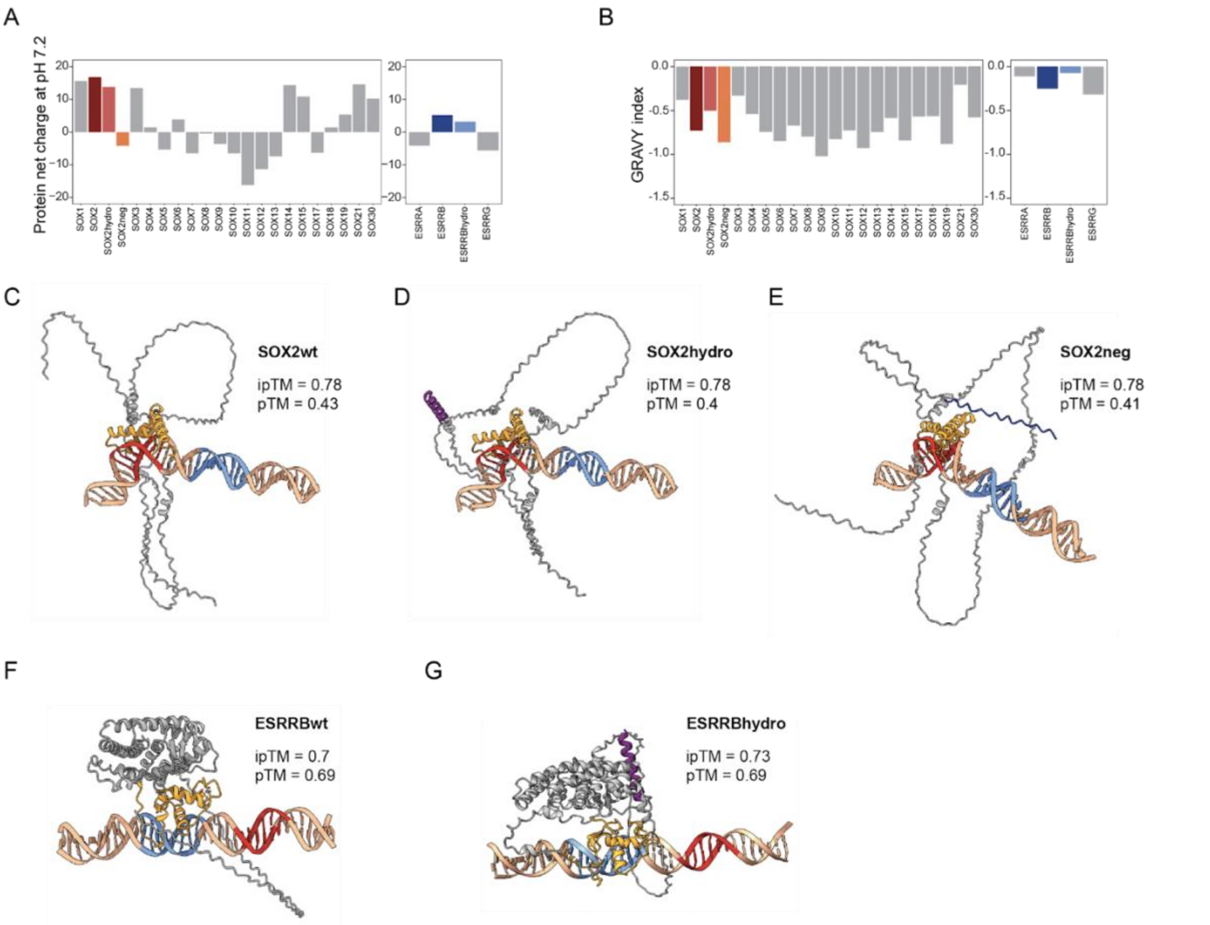


**Figure S1 - related to Figure 1. In silico characterization of vTFs. A-B**) Protein net-charge at pH 7.2 (**A**) and GRAVY index of hydrophobicity (**B**) of mouse TFs belonging to the SOX family (left) and ERR family (right), plus SOX2hydro, SOX2neg and ESRRBhydro variants. **C-G**) Alphafold 3 predictions of SOX2wt (**C**), SOX2hydro (**D**), SOX2neg (**E**), ESRRBwt (**F**) and ESRRBhydro (**G**) folding and binding to a DNA sequence containing both a SOX2 (red) and ESRRB (blue) binding site [63], and corresponding predicted template modeling (pTM) and interface predicted template modeling (iPTM) scores. Purple depicts the hydrophobic amino acid variant, blue the negative charges.

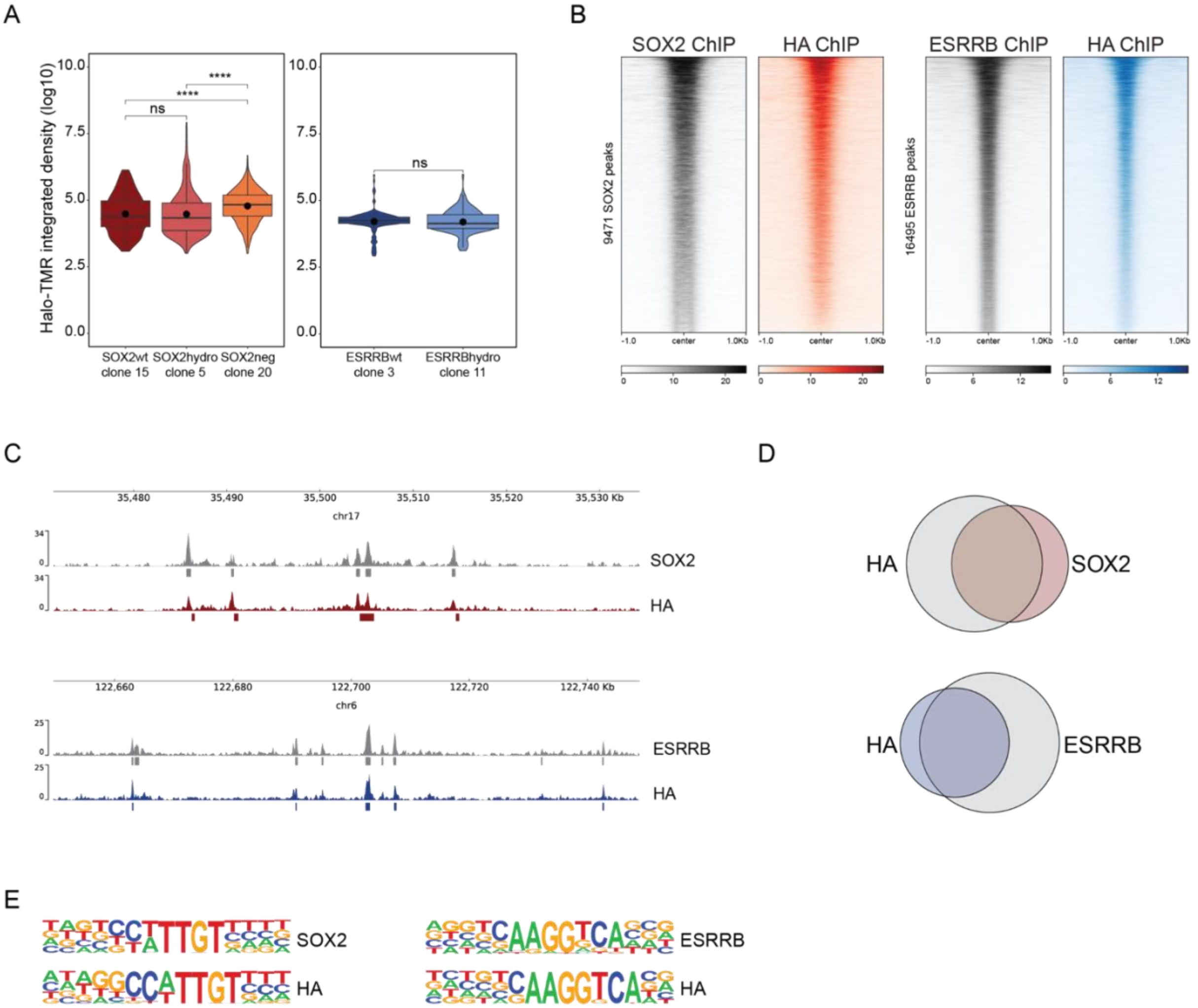


**Figure S2 - related to Figure 1. Validation of vTF-Halo expressing mESCs. A**) Quantification of vTF expression in selected clones, measured by live imaging of Halo-stained cells. P-value: ns, non significative; ****, p<0.0001, Wilcoxon test. n = at least 120 cells in two replicates. **B**) Validation of wild-type (wt) SOX2wt (left) and ESRRBwt (right) fusion proteins genomic binding by ChIP-Seq. (left) Enrichment of endogenous SOX2 (grey) and SOX2wt-Halo-HA (red) at SOX2 peaks. (right) Enrichment of endogenous ESRRB (grey) and ESRRBwt-Halo-HA (blue) at ESRRB peaks. **C-D**) Genome tracks (**C**) and overlap of detected peaks (**D**) derived from ChIP-Seq performed using an anti-SOX2 or anti-HA antibody in cells expressing the Halo-SOX2wt-HA construct (top) and or an anti-ESRRB or anti-HA antibody in cells expressing the ESRRBwt-Halo-HA construct (bottom). In **C**, squares represent called peaks. **E**) Top motif detected by Homer de novo motifs analysis (see Methods) at detected peaks.

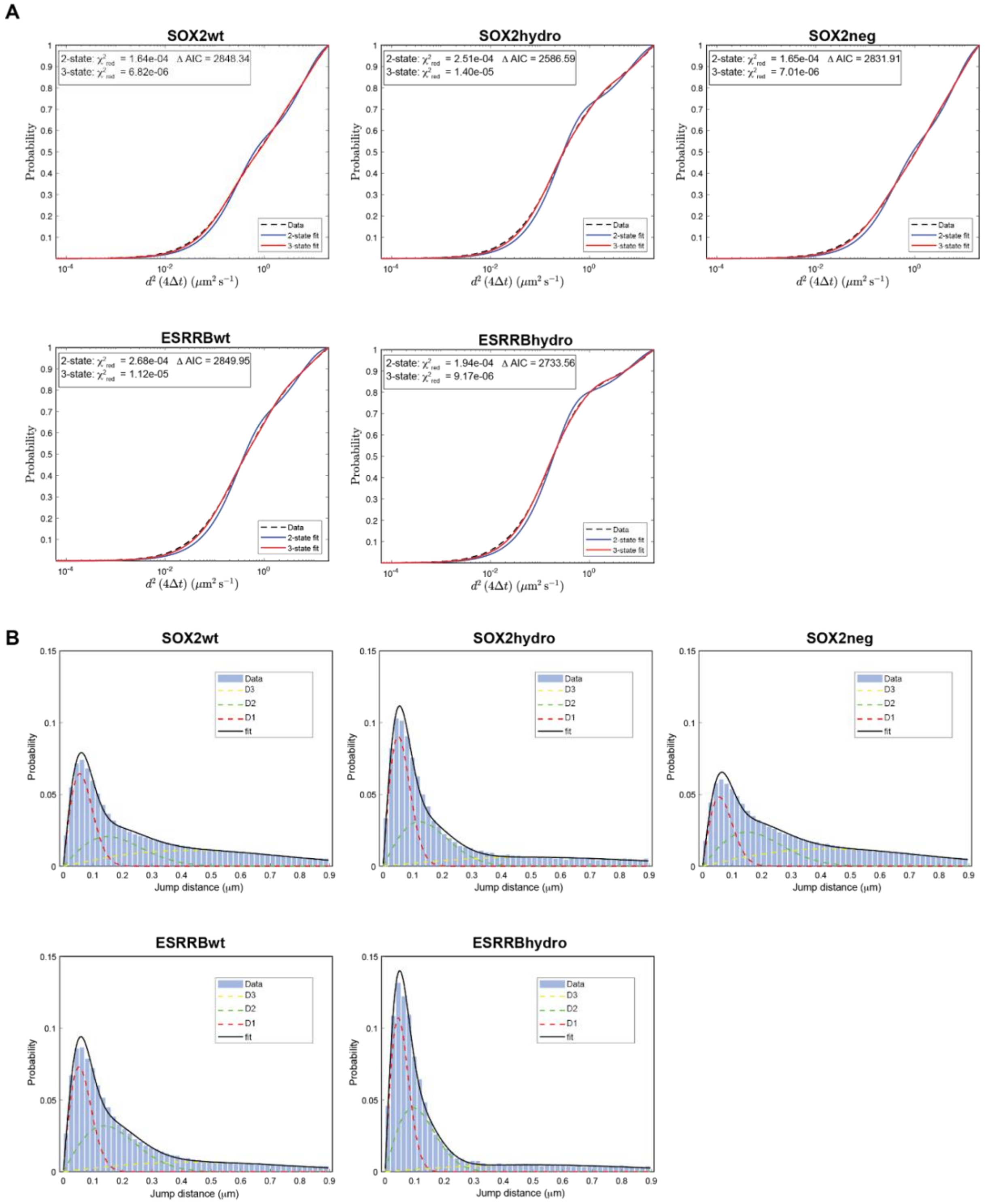


**Figure S3 - related to Figure 2. Analysis of diffusion measurements. A**) Cumulative distributions of jump distances (black dashed lines) derived from analysis of continuous movies and fits with a two-state (blue) or three-state (red) diffusion model. Data are best described by a three-states fit as determined by the reduced $\chi^2$ analysis and the Akaike Information Criterion (AIC). **B**) Jump distance distributions (blue histograms) of continuous movies fit with a three component diffusion model (black) and the single components (red, green and yellow).

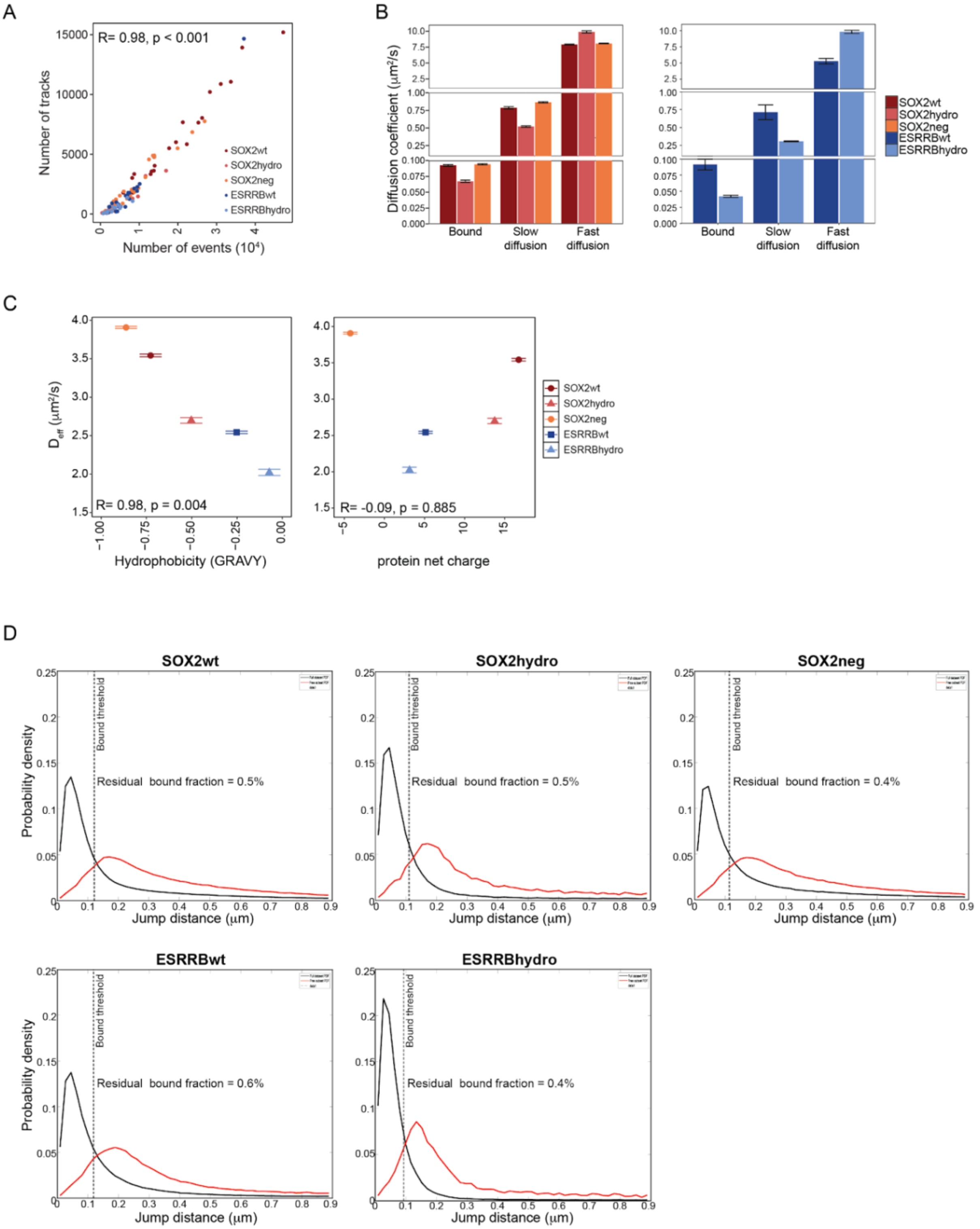


**Figure S4 - related to Figure 2**. **Diffusion coefficients and residual bound fractions. A**) Number of single-molecule tracks plotted against the number of all events (spots detected per frame) from continuous movies. **B**) Diffusion coefficients of bound, slow and fast diffusing molecules, derived from fitting the continuous movies with a three-states diffusion model (Methods and Supplementary Table 3). Data are provided as mean values ± standard deviation of 500 resamplings with 80% of data. **C**) Correlation between the average diffusion coefficient Deff and TF hydrophobicity (as measured by GRAVY index, left) or net charge (right). **D**) Jump distance probability density function (PDFs) for the full datasets (black) and free subset (red). The dashed line marks the bound-state threshold, calculated from the fitted bound diffusion coefficient (D1) and frame interval (Δt). The residual bound fraction in the free trajectories pool was obtained by integrating the free subset PDF below this threshold and normalizing to the bound fraction of the full dataset.

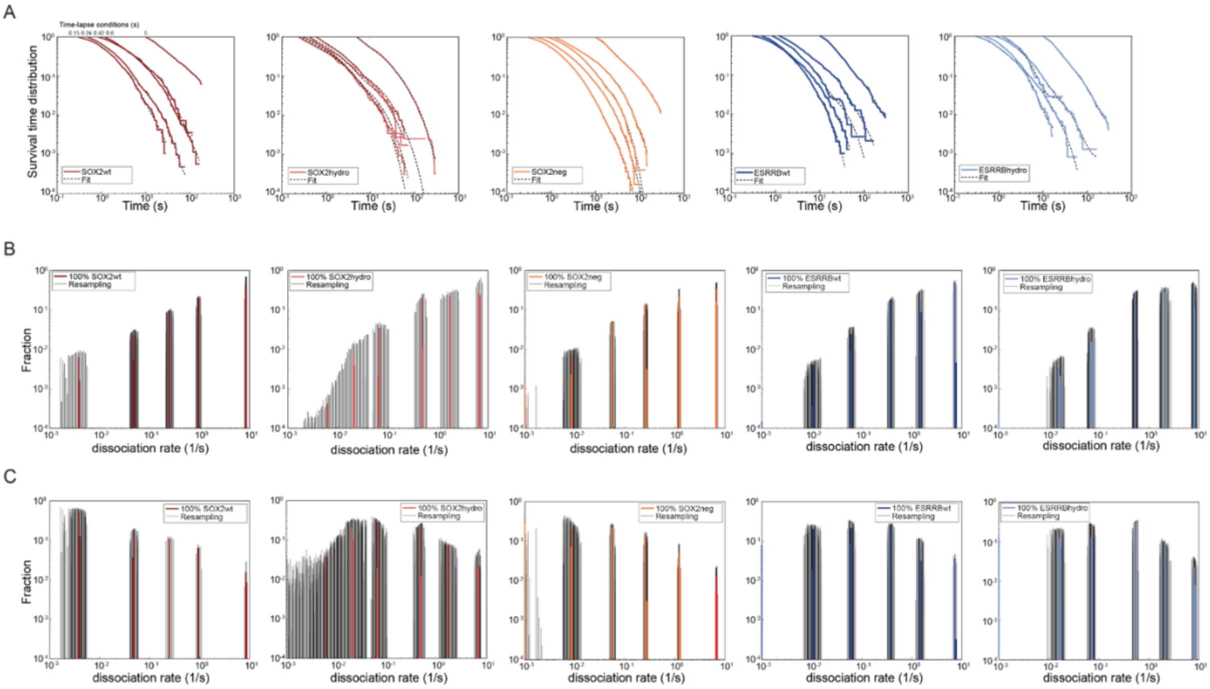


**Figure S5 - related to Figure 3. Survival time distributions and raw GRID data. A**) Survival time distributions (coloured line) and GRID fit (dashed black line) of wt and vTFs. Time-lapse conditions are indicated on top. **B-C**) Event (**B**) and state (**C**) spectra of wt and vTFs obtained with GRID analysis (see Methods). Resampling was performed by repeating the GRID analysis 500 times with 80% of randomly selected survival time distributions. Statistics for time-lapse imaging experiments are provided in Supplementary Table 4.

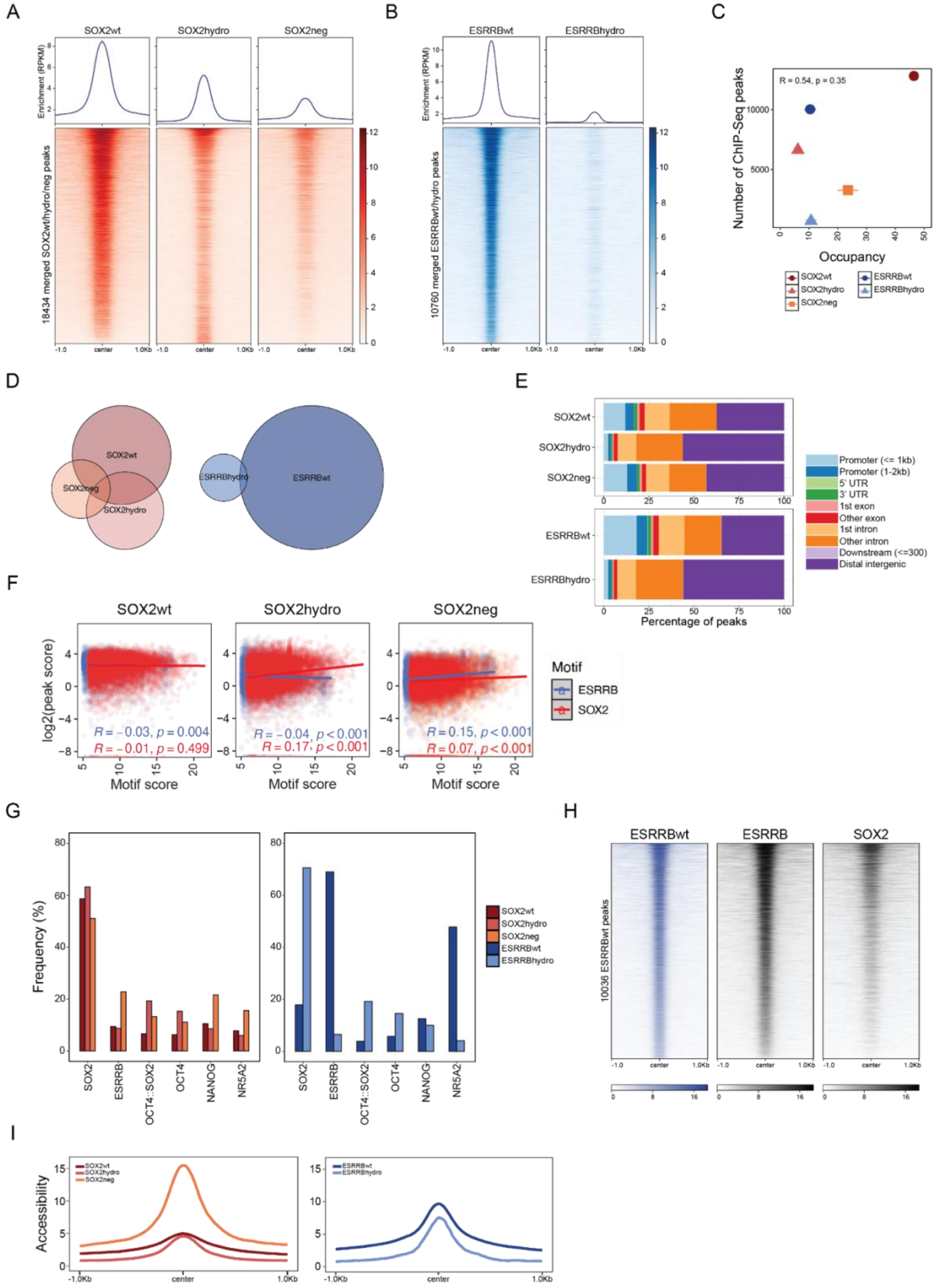


**Figure S6 - related to Figure 4. Characterization of vTFs genomic binding A**) Heatmap of ChIP-Seq enrichment of SOX2wt, SOX2hydro and SOX2neg at SOX2 peaks. **B**) Heatmap of ChIP-Seq enrichment of ESRRBwt and ESRRBhydro at ESRRB peaks. **C**) Number of peaks called by ChIP-Seq of each TF variant plotted against their respective occupancy. **D**) Overlap of regions bound by SOX2wt, SOX2hydro and SOX2neg (left) and ESRRBwt and ESRRBhydro (right). **E**) Percentage of representation of each genomic feature (see color legend) in each vTF peak set. **F**) ChIP-Seq enrichment peak score (log2 RPKM) for SOX2wt, SOX2hydro and SOX2neg plotted against the ESRRB or SOX2 motif quality score for the same peak. **G**) Frequency of SOX2, ESRRB, OCT4::SOX2, OCT4, NANOG and NR5A2 motifs at vTFs called peaks (see Methods). **H**) Heatmap of ChIP-Seq enrichment of ESRRBwt, endogenous ESRRB and endogenous SOX2 at ESRRBwt peaks. **I**) Enrichment profile of ATAC-Seq signal at vTF bound regions.

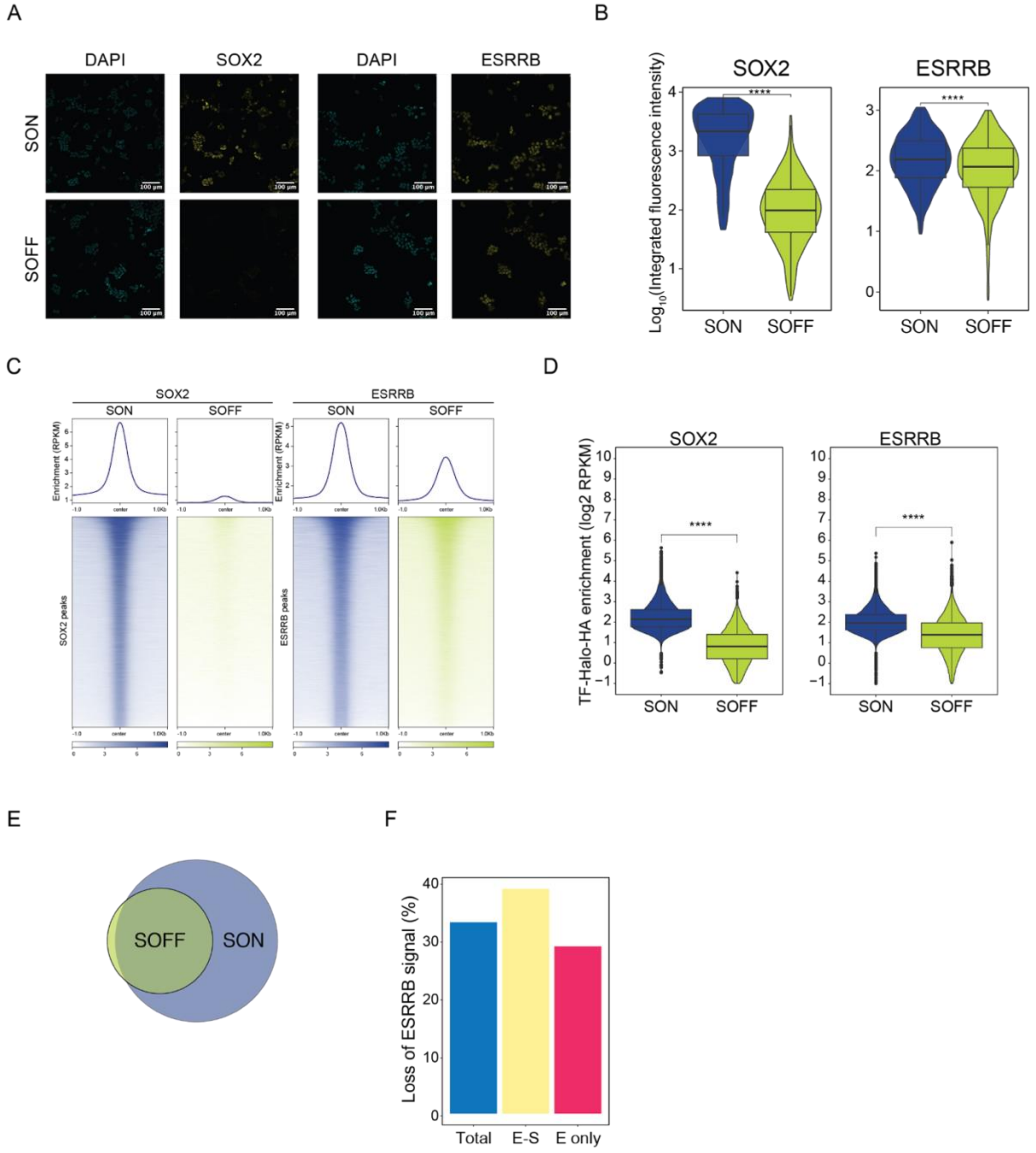


**Figure S7 - related to Figure 5**. **ESRRB expression and binding in 2TS22C cells. A**) Microscopy images of SOX2 and ESRRB immunofluorescence in 2TS22C cells in control conditions (SON) and upon 26h of dox treatment (SOFF). Nuclei were additionally stained with DAPI. **B**) Quantification of SOX2 and ESRRB signals in immunofluorescence experiments (N = at least 800 cells) in control conditions (SON) and upon 26h of dox treatment (SOFF). **C**) Heatmap of ChIP-Seq enrichment of endogenous SOX2 (left) and ESRRB (right). **D**) Average enrichment score (log2 RPKM) of endogenous SOX2 (left) and ESRRB (right) ChIP-Seq in presence and absence of SOX2. **E**) Overlap of endogenous ESRRB bound regions (as called peaks) in presence and absence of SOX2. **F**) Percentage of ESRRB ChIP-Seq score reduction in SOFF condition compared to SON at all called peaks (Total), SOX2-cobound regions (E-S) and regions not bound by SOX2 in SON (E only).

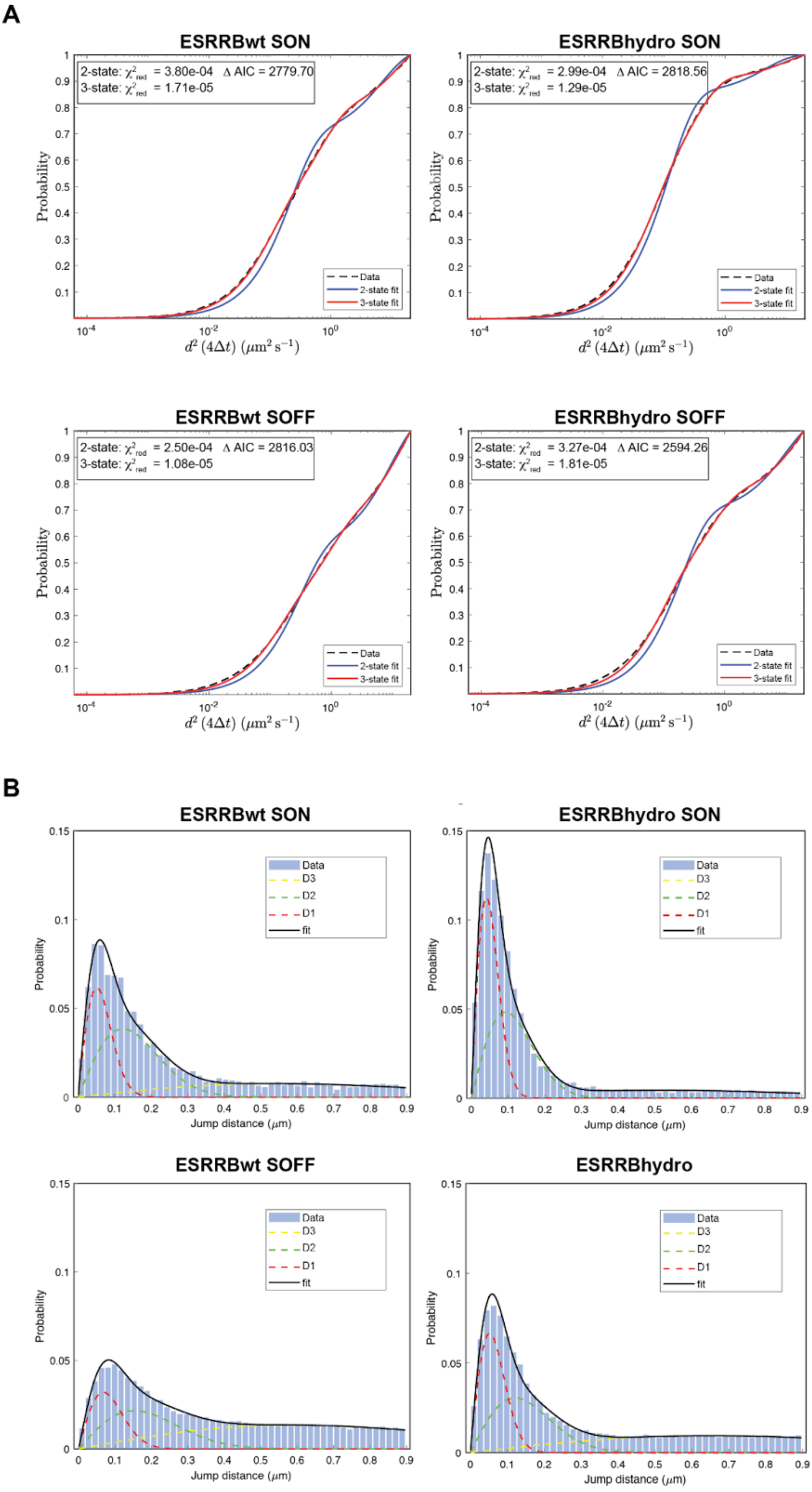
A
ESRRBwt SON
2-state: $\chi^2_{red}$ = 3.80e-04 Δ AIC = 2779.70
3-state: $\chi^2_{red}$ = 1.71e-05
ESRRBhydro SON
2-state: $\chi^2_{red}$ = 2.99e-04 Δ AIC = 2818.56
3-state: $\chi^2_{red}$ = 1.29e-05
ESRRBwt SOFF
2-state: $\chi^2_{red}$ = 2.50e-04 Δ AIC = 2816.03
3-state: $\chi^2_{red}$ = 1.08e-05
ESRRBhydro SOFF
2-state: $\chi^2_{red}$ = 3.27e-04 Δ AIC = 2594.26
3-state: $\chi^2_{red}$ = 1.81e-05
Data
2-state fit
3-state fit
Probability
$d^2\,(4\Delta t)\ (\mu m^2\,s^{-1})$
B
ESRRBwt SON
ESRRBhydro SON
ESRRBwt SOFF
ESRRBhydro
Data
D3
D2
D1
fit
Probability
Jump distance ($\mu$m)

**Figure S8 - related to Figure 5. Analysis of diffusion measurements. A**) Cumulative distributions of jump distances (black dashed lines) derived from analysis of continuous movies of 2TS22C cells and fits with a two-state (blue) or three-state (red) diffusion model. Data are best described by a three-states fit as determined by the reduced $\chi^2$ analysis and the Akaike Information Criterion (AIC). **B**) Jump distance distributions (blue histograms) of continuous movies fit with a three component diffusion model (black) and the single components (red, green and yellow).

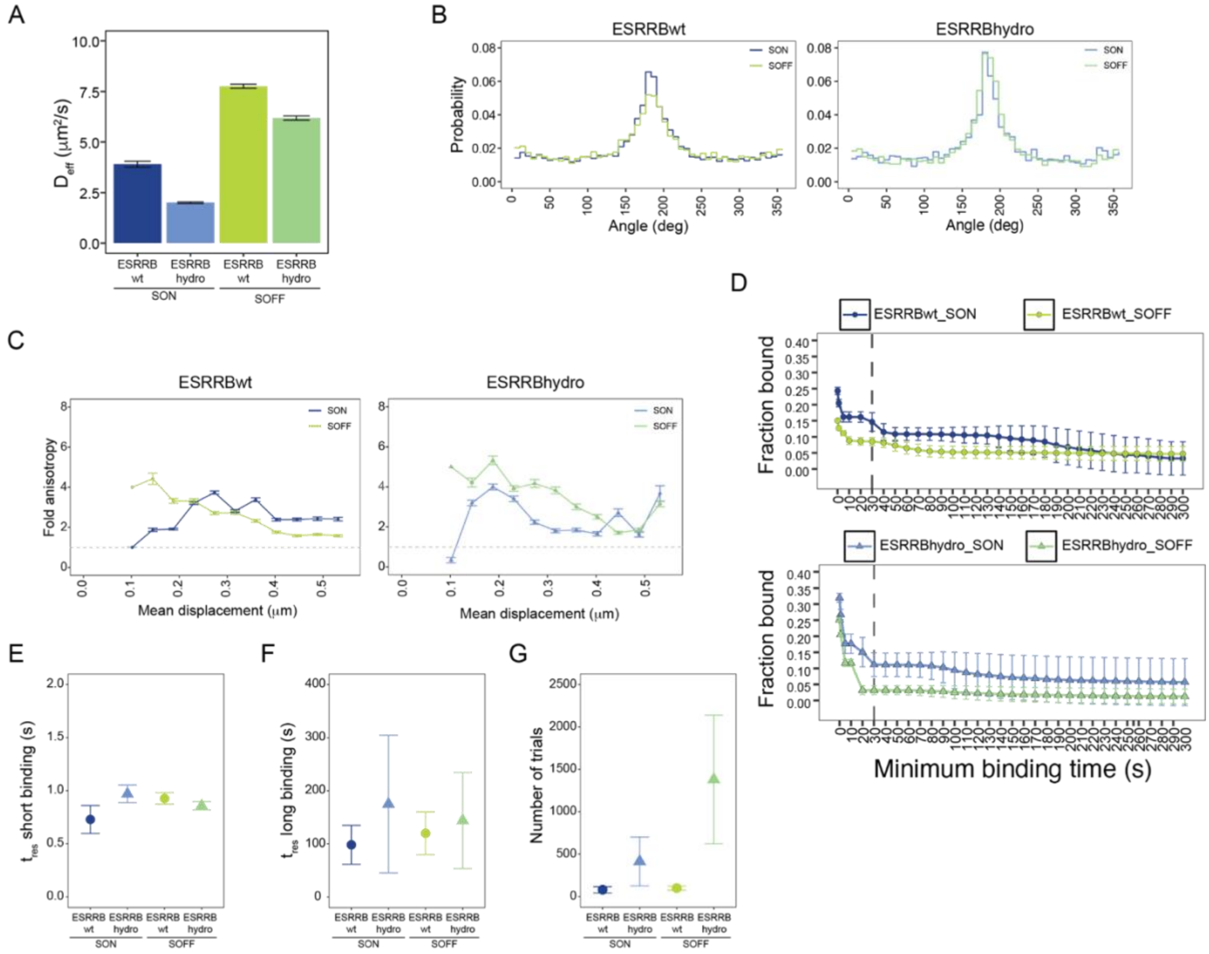


**Figure S9 - related to Figure 5. Additional ESRRB search parameters in presence and absence of SOX2. A**) Average diffusion coefficient $D_{eff}$ as measure of overall TF mobility, derived from bound fractions in Figure 5b and diffusion coefficients in Figure S9a (Methods and Supplementary Table 3). Data are provided as mean values ± standard deviation of 500 resamplings with 80% of data. **B**) Histograms of angle distribution probability for ESRRBwt and ESRRBhydro in SON and SOFF conditions. **C**) Fold anisotropy ratio $f_{180/0}$ plotted as function of mean displacement length over all lag times. Data are provided as mean values ± SD of 500 resamplings with 80% of data. **D**) Cumulative binding curves showing the proportion of molecules that remain bound for at least the indicated residence time. Curves were obtained by weighing the total bound fraction by GRID dissociation rate amplitudes and plotting it as a function of residence time (see Methods). At binding time 0 s, the corresponding Y value corresponds to the total bound fraction determined by continuous diffusion movies (Figure 5b). Data are provided as mean values ± standard deviation of 500 resamplings with 80% of data. **E-F**) Average residence times of all binding events lasting less than 30 s (**E**) and more than 30 s (**F**). Data are provided as mean values ± standard deviation. **G**) Number of unspecific encounters determined for each TFs from the amplitude of the event spectrum above 30s (Methods and Supplementary Table 7), shown as mean values ± error from error propagation.

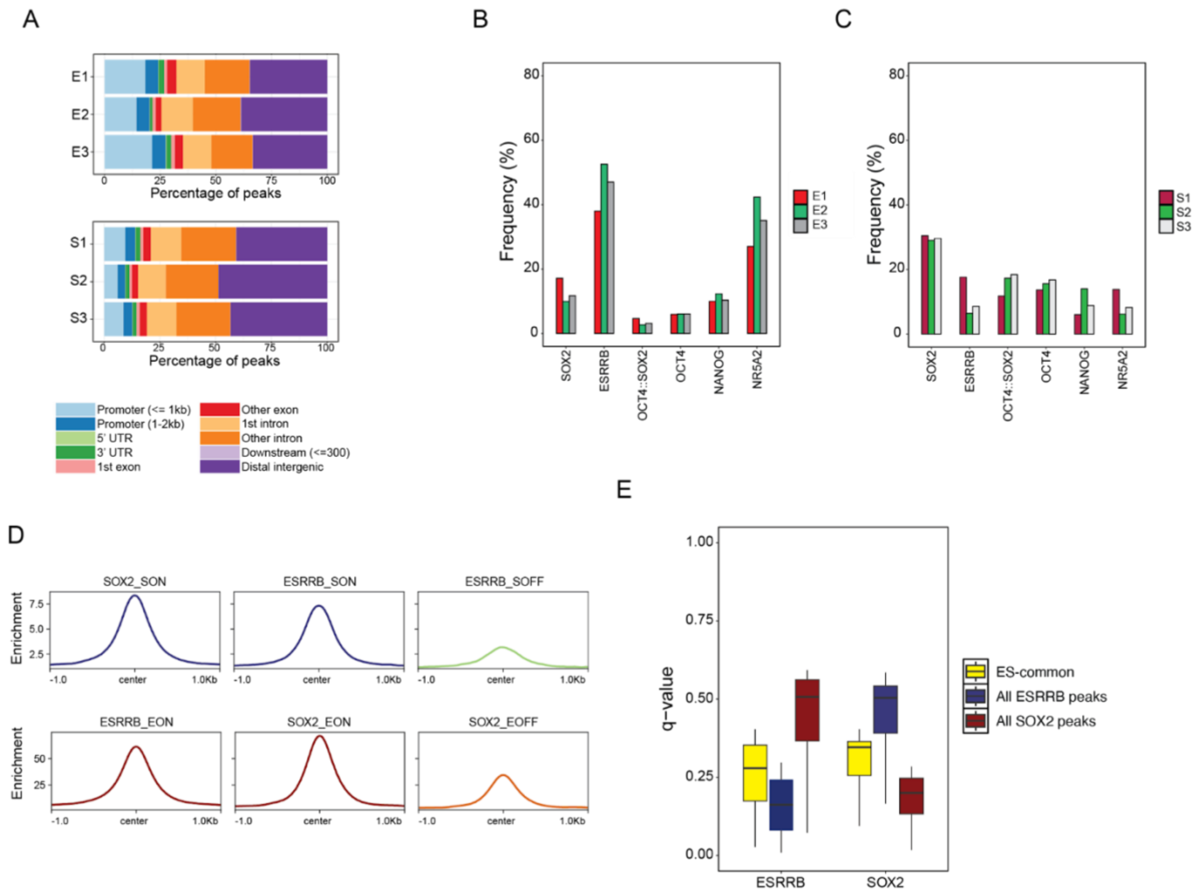


**Figure S10 - related to figure 6. Characterization of cluster regions A**) Percentage of representation of each genomic feature (see color legend) in each cluster defined in Fig.6A. **B-C**) Frequency of SOX2, ESRRB, OCT4::SOX2, OCT4, NANOG and NR5A2 motifs at called peaks in each cluster. **D**) Enrichment profile of SOX2 and ESRRB in 2TS22C (top) and EKOiE from published data (bottom) at ES-dependent sites (see Figure 6D). **E**) Quality of ESRRB and SOX2 motifs at all SOX2 and ESRRB sites and at ES-dependent regions (see Methods).

### Supplementary tables

Supplementary Table 1. List of primers used in this study

| Sequence | Primer ID |
|---|---|
| TACATGgtcgacATGGGAatcgatTCGTCCGAAGACAGGCAC | ESRRB_cDNA_SalI_ClaI Fw |
| CATGTAtctagaCACCTTGGCCTCCAGCAT | ESRRB_cDNA_XbaI Rv |
| TACATGtctagaGCAGAAATCGGTACTGGCTTTC | Halo_XbaI Fw |
| ACTGTAGGCGCGCCTCAAGCGTAATCTGGAACATCGTATGGGTACATAGGAGCATAATCTGGAACATCATAGGGATAAGCGTAATCTGGAACATCGTATGGGTAGTTATCGCTCTGAAAGTACAGATCCT | Halo_3xHA_Stop_AscI Rv |
| TACATGtctagatataacatgatggagacggagctg | SOX2_cDNA_XbaI Fw |
| actgtaGGCGCGCCTCAAGCGTAATCTGGAACATCGTATGGGTACATAGGAGCATAATCTGGAACATCATAGGGATAAGCGTAATCTGGAACATCGTATGGGTAatcgatTCCcatgtgcgacaggggcagt | SOX2_cDNA_ClaI_3xHA_AscI Rv |
| TACATGatcgatCTGATCCTGATCCTGATCCTGATCCTGATCCTGATCCTGATCCTGATCCTGATCatcgatTACATG | ESRRBhydro Fw |
| CATGTAatcgatGATCAGGATCAGGATCAGGATCAGGATCAGGATCAGGATCAGGATCAGGATCAGatcgatCATGTA | ESRRBhydro Rv |
| TACATGatcgatCTGTTCCTGTTCCTGTTCCTGTTCCTGTTCCTGTTCCTGTTCCTGTTCCTGTTCatcgatTACATG | SOX2hydro Fw |
| CATGTAatcgatGAACAGGAACAGGAACAGGAACAGGAACAGGAACAGGAACAGGAACAGGAACAGatcgatCATGTA | SOX2hydro Rv |
| TACATGatcgatGACGAGGACGAGGACGAGGACGAGGACGAGGACGAGGACGAGGACGAGGACGAGatcgatTACATG | SOX2neg Fw |
| CATGTAatcgatCTCGTCCTCGTCCTCGTCCTCGTCCTCGTCCTCGTCCTCGTCCTCGTCCTCGTCatcgatCATGTA | SOX2neg Rv |
| GGTGTCGTGAGGATCATGGGAATCGATTCGTCCG | Infusion ESRRBwt-Halo-HA in pLV-EF1a plasmid Fw |
| GGTGTCGTGAGGATCATGGGAATCGATCTGATCCTGATCC | Infusion ESRRBhydro-Halo-HA in pLV-EF1a plasmid Fw |
| TCGAGCGGCCGCCACAGCGTAATCTGGAACATCGTATGG | Infusion ESRRBwt/hydro-Halo-HA in pLV-EF1a plasmid Rv |

Supplementary table 2. Statistics of continuous movies and tracking statistics .

| TF | # movies | # tracks | # events |
|---|---|---|---|
| SOX2wt | 20 | 131162 | 408236 |
| SOX2hydro | 18 | 13045 | 75003 |
| SOX2neg | 29 | 66631 | 246060 |
| ESRRBwt | 21 | 37465 | 152003 |
| ESRRBhydro | 16 | 6771 | 52464 |
| ESRRBwt SON | 22 | 2611 | 36494 |
| ESRRBwt SOFF | 26 | 21008 | 234115 |
| ESRRBhydro SON | 20 | 6866 | 46611 |
| ESRRBhydro SOFF | 28 | 18171 | 216248 |

Supplementary Table 3. Diffusion coefficients and fractions derived from cumulative jump distance distribution of fast SMT data in Figure S3 and shown in Figure 2 and S4.

| TF | D1 | D2 | D3 | A1 (bound) | A2 (slow) | A3 (fast) |
|---|---|---|---|---|---|---|
| SOX2wt | 0.092 ± 0.001 | 0.786 ± 0.016 | 7.93 ± 0.07 | 0.283 ± 0.002 | 0.304 ± 0.001 | 0.414 ± 0.001 |
| SOX2hydro | 0.067 ± 0.002 | 0.520 ± 0.011 | 9.90 ± 0.18 | 0.334 ± 0.006 | 0.418 ± 0.005 | 0.248 ± 0.002 |
| SOX2neg | 0.094 ± 0.001 | 0.865 ± 0.009 | 8.08 ± 0.04 | 0.217 ± 0.002 | 0.338 ± 0.001 | 0.445 ± 0.001 |
| ESRRBwt | 0.082 ± 0.001 | 0.727 ± 0.007 | 7.95 ± 0.07 | 0.308 ± 0.002 | 0.432 ± 0.002 | 0.261 ± 0.001 |
| ESRRBhydro | 0.04 ± 0.002 | 0.308 ± 0.006 | 9.83 ± 0.23 | 0.314 ± 0.008 | 0.497 ± 0.008 | 0.189 ± 0.002 |
| ESRRBwt SON | 0.068 ± 0.004 | 0.595 ± 0.02 | 12.34 ± 0.58 | 0.243 ± 0.011 | 0.464 ± 0.009 | 0.293 ± 0.005 |
| ESRRBwt SOFF | 0.112 ± 0.005 | 0.867 ± 0.023 | 14.39 ± 0.24 | 0.15 ± 0.005 | 0.33 ± 0.004 | 0.518 ± 0.003 |
| ESRRBhydro SON | 0.036 ± 0.001 | 0.304 ± 0.006 | 10.49 ± 0.26 | 0.319 ± 0.014 | 0.505 ± 0.013 | 0.175 ± 0.003 |
| ESRRBhydro SOFF | 0.069 ± 0.002 | 0.537 ± 0.01 | 16.68 ± 0.34 | 0.251 ± 0.005 | 0.392 ± 0.004 | 0.357 ± 0.002 |

Supplementary table 4. Statistics of time-lapse movies and tracking.

| Time lapse conditions | | SOX2wt | SOX2hydro | SOX2neg | ESRRBwt | ESRRBhydro |
|---|---|---|---|---|---|---|
| 150ms | # movies | 15 | 14 | 8 | 8 | 4 |
| | # tracks | 1192 | 4764 | 11941 | 2273 | 516 |
| | # all events | 11048 | 21624 | 137230 | 21505 | 3868 |
| 240ms | # movies | 16 | 4 | 7 | 11 | 6 |
| | # tracks | 3093 | 530 | 8893 | 1317 | 1866 |
| | # all events | 24525 | 2552 | 87803 | 8001 | 9371 |
| 420ms | # movies | 7 | 4 | 9 | 8 | 6 |
| | # tracks | 2263 | 299 | 3804 | 655 | 1166 |
| | # all events | 14219 | 1682 | 34364 | 4504 | 5360 |
| 600ms | # movies | 15 | 5 | 7 | 7 | 4 |
| | # tracks | 490 | 362 | 2930 | 693 | 68 |
| | # all events | 3787 | 1892 | 25726 | 5714 | 626 |
| 5.06sec | # movies | 12 | 17 | 10 | 12 | 21 |
| | # tracks | 2829 | 6199 | 5361 | 2746 | 4499 |
| | # all events | 15855 | 28090 | 30132 | 18764 | 25859 |
| Tot tracks | | 9867 | 12154 | 32929 | 7684 | 8155 |

Supplementary Table 5. Search parameters of vTFs as shown in Figure 2 and S4.

| TF | $pb_s$ | $pb_u$ | $k_{off,s}$ | $k_{off,u}$ | $N_{trials}$ | $t_{3D}$ | $T_{search}$ |
|---|---|---|---|---|---|---|---|
| SOX2wt | 0.18 ± 0.007 | 0.10 ± 0.007 | 0.004 ± 0.0005 | 0.81 ± 0.04 | 124 ± 9.6 | 8.55 ± 0.71 | 1217 ± 130 |
| SOX2hydro | 0.11 ± 0.017 | 0.22 ± 0.017 | 0.019 ± 0.002 | 0.82 ± 0.08 | 82 ± 22 | 3.68 ± 0.44 | 400 ± 115 |
| SOX2neg | 0.12 ± 0.009 | 0.09 ± 0.009 | 0.005 ± 0.0006 | 0.58 ± 0.015 | 95 ± 8.45 | 13.8 ± 1.33 | 1471 ± 184 |
| ESRRBwt | 0.086 ± 0.011 | 0.199 ± 0.011 | 0.0083 ± 0.0012 | 0.767 ± 0.035 | 216 ± 30 | 4.66 ± 0.34 | 1286 ± 197 |
| ESRRBhydro | 0.103 ± 0.012 | 0.21 ± 0.013 | 0.0097 ± 0.0012 | 0.873 ± 0.046 | 183 ± 28 | 3.74 ± 0.30 | 893 ± 146 |

Supplementary table 6. Statistics of time-lapse movies and tracking in 2TS22C cells.

| **Time lapse conditions** | | **ESRRBwt SON** | **ESRRBwt SOFF** | **ESRRBhydro SON** | **ESRRBhydro SOFF** |
|---|---|---|---|---|---|
| 150ms | # movies | 7 | 16 | 11 | 13 |
| | # tracks | 2023 | 2615 | 637 | 2265 |
| | # all events | 26230 | 76224 | 19013 | 78612 |
| 240ms | # movies | 8 | 12 | 9 | 12 |
| | # tracks | 604 | 2970 | 1126 | 1858 |
| | # all events | 13087 | 48587 | 15881 | 29264 |
| 420ms | # movies | 11 | 7 | 7 | 9 |
| | # tracks | 1127 | 505 | 752 | 985 |
| | # all events | 13439 | 10132 | 10366 | 14191 |
| 600ms | # movies | 4 | 16 | 5 | 10 |
| | # tracks | 179 | 1914 | 306 | 2610 |
| | # all events | 3100 | 17302 | 3403 | 16119 |
| 5.06sec | # movies | 7 | 8 | 9 | 5 |
| | # tracks | 109 | 383 | 104 | 236 |
| | # all events | 594 | 2644 | 907 | 1213 |
| Tot tracks | | 4042 | 8387 | 2925 | 7954 |

Supplementary Table 7. Search parameters of vTFs as shown in Figure 2 and S4.

| **TF** | **$pb_s$** | **$pb_u$** | **$k_{off,s}$** | **$k_{off,u}$** | **$N_{trials}$** | **$t_{3D}$** | **$T_{search}$** |
|---|---|---|---|---|---|---|---|
| ESRRBwt SON | 0.147 ± 0.029 | 0.096 ± 0.028 | 0.01 ± 0.004 | 1.37 ± 0.25 | 81 ± 37 | 5.7 ± 1.93 | 519.9 ± 287 |
| ESRRBwt SOFF | 0.085 ± 0.012 | 0.065 ± 0.011 | 0.008 ± 0.003 | 1.08 ± 0.06 | 100 ± 24 | 12.12 ± 2.3 | 1308 ± 389 |
| ESRRBhydro SON | 0.111 ± 0.037 | 0.21 ± 0.034 | 0.006 ± 0.004 | 1.03 ± 0.09 | 414 ± 288 | 3.12 ± 0.63 | 1717 ± 1224 |
| ESRRBhydro SOFF | 0.032 ± 0.014 | 0.22 ± 0.014 | 0.007 ± 0.004 | 1.16 ± 0.05 | 1380 ± 758 | 2.94 ± 0.23 | 5249 ± 2903 |
